# Autism-like behavior induced by conditional ablation of the *Bassoon* gene in GABAergic interneurons

**DOI:** 10.1101/2025.11.20.689526

**Authors:** Anil Annamneedi, Carolina Montenegro-Venegas, Gürsel Çalışkan, Endre Marosi, Armand Blondiaux, Nikhil Tiwari, Misbah Razzaq, Judith C. Kreutzmann, Lucie P. Pellissier, Anna Fejtova, Sanja Mikulovic, Markus Fendt, Eckart D. Gundelfinger, Oliver Stork

## Abstract

Synaptic dysfunction and resulting imbalance of excitatory vs. inhibitory transmission are fundamental aspects of autism spectrum disorders (ASD). Here we addressed the role of inhibitory synapse disturbances in ASD employing mice with a conditional ablation in inhibitory forebrain interneurons of the presynaptic active zone scaffolding protein Bassoon (Bsn), a key factor in synaptic development, activity and maintenance. The conditional *Bsn* gene knock out resulted in a reduction of synaptic vesicles and reduced synaptic efficacy of affected inhibitory synapses as well as diminished GABAergic inhibition in hippocampal pyramidal neurons. *Bsn* mutants further displayed a reconfiguration of the hippocampal synaptic network *in vivo*, as seen in subfield-specific changes of inhibitory synaptic markers and reduced numbers of parvalbumin interneurons, culminating in disturbance of hippocampal network activity patterns and profound mitochondrial and metabolic dysregulation. Importantly, the mutant mice developed behavioral abnormalities reminiscent of ASD including widespread social behavioral deficits and novelty-induced hyperarousal with altered motor behavior, increased anxiety and epileptiform activity. *Bsn* conditional knock out mice thus provide strong evidence for a causal involvement of GABAergic synapses in the emergence of ASD-related behavioral and physiological phenotypes.

## Introduction

Bassoon is a vertebrate-specific protein critically involved in synaptic development, maintenance, and function. As a key organizer of the presynaptic active zone of neurotransmitter release [1], it is found in both excitatory and inhibitory presynapses, as well as neuromodulatory varicosities [2, 3]. Bassoon mediates active zone formation at nascent synapses during development and controls the localization of voltage-gated calcium (Ca^2+^) channels at the presynapse, regulating neurotransmission [4–7]. Further studies have demonstrated its role in presynaptic proteome regulation via autophagy and proteasomal degradation pathways [8–10] and in synapse-to-nucleus communication [11].

Mutations in the human *BSN* gene have been associated with neurodevelopmental and neuropsychiatric conditions. These include autism spectrum disorders (ASD) [12, 13], intellectual disability [14], attention deficit hyperactivity disorder (ADHD) [12], schizophrenia [15], as well as juvenile epilepsies, e.g., Landau-Kleffner syndrome and febrile seizures [12, 16–18]. In mice, deletion of Bassoon impairs the assembly, maintenance, integrity, and function of presynaptic boutons in glutamatergic synapses and leads to epileptic seizures and sensory impairments [5, 19–23]. The severe phenotype of these animals precludes a more fine-grained analysis of prospective neurobehavioral deficits [19, 24–26]. In a conditional mouse mutant deficient for *Bsn* expression in glutamatergic forebrain neurons (*Bsn^Emx1^*cKO), we confirmed the relevance of Bassoon for hippocampal circuit development and maturation of the dentate gyrus (DG) as well as epilepsy, but behavioral features reminiscent of ASD or intellectual disability were not detected [26–28].

Therefore, and considering the importance of inhibitory synapse function in ASD [29, 30] we established a mouse line lacking *Bsn* expression in GABAergic interneurons (*Bsn^Dlx5/6^* cKO mice). To avoid putatively detrimental effects complete elimination of interneurons, and considering the particular relevance of parvalbumin (PV) interneurons in ASD [31, 32] we chose a *distal-less Dlx5/6* promoter controlled Cre recombinase, which predominantly target forebrain interneurons that express PV [33]. We investigated the impact of the *Bsn* mutation on GABAergic inhibition in *Bsn^Dlx5/6^*cKO mice at cellular and systems levels and changes in network physiology and describe the emergence of ASD-like behavioral features including comprehensive social deficits, abnormal motor behavior patterns and anxiety-like behavior, as well as metabolic dysregulation in the hippocampus of these animals. Our data shed light on the importance of Bassoon-mediated inhibitory synapse functions in autism spectrum disorders and comorbid conditions.

## Materials and methods

### Animals

Animal experiments performed in this study were conducted in accordance with the European and German regulations for animal experiments. All the experiments were approved by Landesverwaltungsamt Sachsen-Anhalt (Az. 42502-2-1303 LIN, 42502-2-1484 LIN, 42502-2-1797 LIN, and 42502-2-1309 UniMD, 42502-2-1676 UniMD). *Tg(dlx5a-cre)1Mekk* driver line (https://www.informatics.jax.org/allele/MGI:3758328) expressing Cre recombinase under dlx6a (distal-less homeobox gene 6a) promoter with an enhancer element (I56i and I56ii) of zebrafish dlx5a and dlx6a open reading frame were used to drive the recombination in GABAergic forebrain neurons [34, 35]. These mice were bred with *Bsn^lx/lx^* mice (*Bsn^tm1.1Arte^*; https://www.informatics.jax.org/allele/-MGI:7484796), carrying a *loxP*-flanked exon 2 of the *Bsn* gene [27] to generate *Bsn^Dlx5/6^*cKO animals, which lack *Bsn* expression in large fraction (about 40%) of inhibitory interneurons [26].

Two to four months-old male *Bsn^Dlx5/6^*cKO mice and control littermates (*Bsn^lx/lx^*) that are considered as wild types (WT) were used for experiments performed in this study. As additional controls, a group of *Bsn^+/+^X Dlx5/6 Cre* (*WT+Dlx5/6 Cre*) *and their Bsn^+/+^* (*WT w/o Cre*) littermates were tested behaviorally (Fig. S10).

### Analysis of GABAergic synapses

For immunocytochemistry and live-cell assays, primary hippocampal cultures from newborn *Bsn^Dlx5/6^*cKO and their WT littermates were prepared, and Synaptotagmin1 luminal domain antibody (Syt1Ab) uptake assays were performed as previously described [10, 36]. Bassoon-containing (Bsn+) were discriminated from Bassoon-lacking (Bsn-) inhibitory presynaptic boutons in *Bsn^Dlx5/6^*cKO hippocampal culture by immunostaining after the uptake experiments. Densities of excitatory and inhibitory synapses were measured 10-30 µm and 60-80 µm away from the cell body, using triple staining for Synaptophysin1, VGLUT1, Shank2 and, Synapthophysin1, Gephyrin, VGAT respectively. For immunohistochemical analysis mice were perfused with PBS, then with 4% PFA and postfixed overnight in same fixative. Brains were cryoprotected in sucrose solution and stored at −80^0^C until used for immunohistochemical staining [27]. A list of antibodies and conditions is provided in the supplementary information.

### Slice electrophysiology

Acute horizontal brain slices including ventral-to-middle hippocampus were prepared, and whole cell patch clamp recordings were performed to measure synaptic potentials as detailed in the supplementary information. Slice preparation for extracellular field recordings were performed similar to our previous studies [27, 37, 38] and synaptic transmission, plasticity and gamma-oscillations were measured as detailed in the supplementary information.

### Behavioral experiments

One batch of male *Bsn^Dlx5/6^* cKO and control mice was analyzed in a battery of behavioral experiments including home cage activity monitoring, open field exploration, elevated plus maze, light-dark test, social recognition and memory, and nest building as described previously with minor modifications [27, 39, 40]. A second batch of mice was studied for their response to female urine and females in estrus [41, 42]. Further details are provided in the supplementary information.

### Protein analysis

For immunoblot analysis hippocampal tissue was dissected, and crude (P2) membrane fraction was prepared as described previously [19]. P2 membrane fractions from both genotypes (N=4 per genotype) were also used for proteomic analysis performed by EMBL Proteomics Core Facility (Heidelberg) (https://www.embl.org/groups/proteomics/). Obtained data were analyzed using the MouseMine (www.mousemine.org) data bioinformatic tool for gene ontology (GO) enrichment analysis [43].

### Mitochondrial function assay

*Bsn^Dlx5/6^*cKO and control primary hippocampal neurons were seeded in quintuples in a poly-L-lysine-coated 96-well XF cell culture microplate (Seahorse Agilent Technologies) at densities of 30,000 cells. For the mito-stress assay, neurons at 14 DIV were used. The oxygen consumption rate (OCR) of the different mitochondrial kinetics was measured following the supplier’s instructions.

### Statistical analysis

Normality test (Shapiro-Wilk test and/or Levenés test) and equal variance test (in case of slice electrophysiology) were performed before commencing with one-way or two-way ANOVA (repeated measures), paired or unpaired Students *t*-tests. In case the data did not pass the normality test, Mann-Whitney *U*-test was used. For the statistical comparison of gamma power, log transformed data was used to reduce variability. To determine whether the probability of inducing recurrent epileptiform discharges was altered, Fisheŕs exact test was used. All the data are represented as means ± standard errors of the mean (SEM). Probability values of p<0.05 were considered as statistically significant. Sample sizes are provided in figure captions for each experiment (N: Number of mice; n: Number of cells or slices).

## Results

### Bassoon deficiency at inhibitory synapses in hippocampal primary neuronal cultures impairs presynaptic functions

For a first basic characterization, we compared synaptic parameters in hippocampal primary cultures from *Bsn^Dlx5/6^*cKO and WT mice (*Bsn^lx/lx^*) at DIV19-21. Inhibitory GABAergic synapses were identified by overlapping immunostaining for the vesicular inhibitory amino acid transporter VGAT, the inhibitory postsynapse-specific scaffolding protein Gephyrin, and the general synaptic vesicle (SV) marker Synaptophysin1 (Syp1) that is present in both inhibitory and excitatory presynaptic boutons (Fig. 1a). The average number of GABAergic synapses along proximal and distal dendrites, i.e. within the first 10-30 μm to the soma and about 60-80 μm distal from the soma, respectively, was not significantly different between WT and *Bsn^Dlx5/6^*cKO cultures [Fig. 1b; proximal: U=111, Mann-Whitney *U*-test; distal: t(29)=0.753, Student’s *t-test*]. Similarly, the synapse density of excitatory synapses, identified by staining with the vesicular glutamate transporter VGLUT1, the postsynaptic scaffolding protein Shank2 and Syp1, was unchanged between the two cultures (Supplementary Fig. S1a, b; proximal: t(39)=0.697; distal: U=165). Assessment of the intensity of immunosignals revealed a slight, but significant reduction of Gephyrin [Fig. 1c; t(42)=2.371] and a clear reduction of VGAT (Fig. 1d; U=95) immunoreactivity in *Bsn^Dlx5/6^*cKO vs. WT cultures. No major changes were observed at excitatory synapses [Supplementary Fig. S1c,d; Shank2: t(54)=0.262; Vglut1: U=320]. Evidently, the distribution of intensities of VGAT immunoreactivity in *Bsn^Dlx5/6^*cKO inhibitory synapses is bimodal (Fig. 1d). Since only about one-third of GABAergic presynapses in these cultures lack Bassoon [26], we assumed that this bimodality could reflect Bassoon-containing (Bsn+) and Bassoon-lacking (Bsn-) groups of synapses. To test this hypothesis and to evaluate potential functional differences of the two groups of synapses we performed co-staining assays with antibodies against Bassoon and VGAT combined with Synaptotagmin1 antibody (Syt1Ab) uptake assays that report on the SV recycling activity of individual synapses. Under basal conditions (Fig. 1e), i.e. intrinsic activity of the cultures, we observed a significant reduction of VGAT immunosignal intensity in Bsn-synapses as compared to Bsn+ synapses in *Bsn^Dlx5/6^*cKO primary cultures [Fig. 1f; F(2,103)=40.81, one-way ANOVA]. No significant difference was observed between Bsn+ synapses and inhibitory synapses in WT control cultures (Fig. 1f). A similar reduction was observed for Syp1 immunoreactivity in Bsn-synapses, while Syp1 levels in Bsn+ synapses and in WT inhibitory synapses did not differ significantly [Fig. 1g; F(2,104)=45.18]. This indicated a reduction of SVs in Bassoon-deficient inhibitory synapses in hippocampal cultures from *Bsn^Dlx5/6^*cKO mice. Consequently, SV cycling, as indicated by lower Syt1Ab uptake, is reduced in these synapse as compared to Bsn+ inhibitory synapses in the same culture and VGAT-positive synapses in WT cultures [Fig. 1h; F(2,107)=36.59].

**Figure 1.**
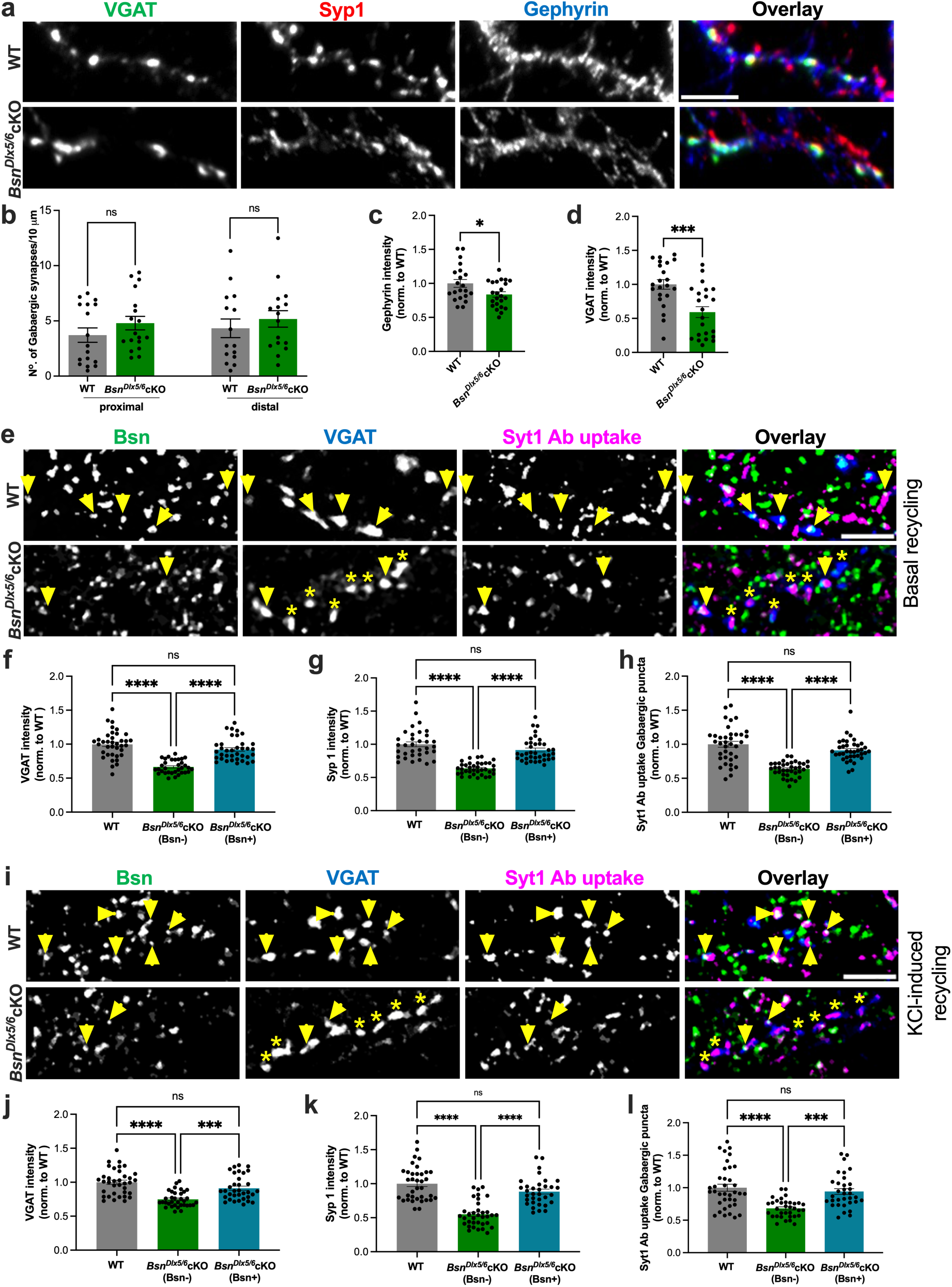
Impaired function of Bassoon-deficient presynapses in *Bsn^Dlx5/6^*cKO hippocampal cultures. **a)** Representative images of 19-21 DIV cultured hippocampal neurons from *WT* and *Bsn^Dlx5/6^cKO* immunostained for VGAT, Syp1 and Gephyrin to label inhibitory synapses. **b)** Quantification of number of inhibitory synapses along 10 μm of proximal and distal dendritic segments revealed no significant change between genotypes. **c,d)** Quantification of normalized immunofluorescence intensity for Gephyrin (**c**) and for VGAT (**d**) revealed decreased levels for both proteins in *Bsn^Dlx5/6^cKO* neurons. Data are obtained from 2 independent cultures per condition. All values are mean ± SEM. Statistical significance was assessed using Student’s *t* test as is depicted in graphs \**P* ≤ 0.05, \*\**P* ≤ 0.01, *** *P* ≤ 0.001. Scale bars, 5 μm. **e)** Representative images of Syt1Ab uptake (magenta) driven by endogenous network activity in hippocampal neurons at 19-21 DIV from *WT* and *Bsn^Dlx5/6^cKO* mice. VGAT (blue) labels the inhibitory presynapses and Bassoon (green) was used to discriminate Bassoon-expressing from Bassoon-deficient presynaptic boutons in *Bsn^Dlx5/6^*cKO neurons. **f,g)** Quantification of immunofluorescence intensity for VGAT (**f**) and Syp1 (**g**) revealed a reduction in the synaptic levels of both proteins in *Bsn^Dlx5/6^*cKO neurons deficient for Bassoon. **h)** Quantification of normalized immunofluorescence of Syt1Ab upon uptake into inhibitory synapses of experiment shown in panel e. **i)** Representative images of Syt1Ab uptake (magenta) upon KCl-induced depolarization (culture and staining conditions as in panel e). **j,k)** Quantification of immunofluorescence intensity for VGAT (j) and Syp1 (k) of experiments shown in i. **l)** Quantification of normalized IF of Syt1Ab in inhibitory synapses of experiments shown in panel i. Data are obtained from 4 independent culture preparations. All values are mean ± SEM. Statistical significance was assessed using one-way ANOVA with Tukeýs multiple comparison test, as depicted in graphs \**P* ≤ 0.05, \*\**P* ≤ 0.01, *** *P* ≤ 0.001, **** *P* ≤ 0.0001. Arrowheads indicate examples of Bassoon-positive (Bsn+), asterisks of Bassoon-negative (Bsn-) synapses. Scale bar is 5 μm.

Brief chemical depolarization by application of high K^+^ (50 mM for 4 minutes) (Fig. 1i), which is supposed to release the total recycling SV pool (TRP) [44, 45], produced similar results. Also in this case, VGAT [Fig. 1j; F(2,101)=20.91] and Syp1 [Fig. 1k; F(2,101)=40.68] immunoreactivity as well as Syt1Ab uptake were significantly reduced in Bsn-inhibitory synapses, while Bsn+ synapses were similar to GABAergic synapses in WT cultures [Fig. 1l; F(2,101)=15.15]. No difference in Syt1Ab uptake was observed in excitatory (VGLUT1-positive) synapses in either condition [Supplementary Fig S1e-g; basal recycling: t(48)=1.840; Vglut1 intensity: U=271; Supplementary Fig S1h-j; evoked recycling: U=204; Vglut1 intensity: U=224]. In essence, these data confirm that GABAergic synapses of only a fraction of inhibitory interneurons are affected by the *Dlx5/6* element driven cKO.

### *Bsn^Dlx5/6^*cKO mice display subfield-specific changes in inhibitory and excitatory synapse density in hippocampus

Immunohistochemical analysis confirmed the selective absence of Bassoon from a subset of VGAT-positive (VGAT+) presynapses in hippocampus and striatum of *Bsn^Dlx5/6^*cKO mice without significant change in excitatory VGLUT1-positive presynapses (Fig. 2, Supplementary Fig. S2a). This appears reasonable considering that only about 20-30% of forebrain synapses are GABAergic [46–48], and assuming that only one third of these are lacking Bassoon, as in cultures from *Bsn^Dlx5/6^*cKO mice. Accordingly, Bassoon levels were also found slightly (but not significantly) reduced in forebrain extracts of *Bsn^Dlx5/6^*cKO, as determined by quantitative immunoblot analysis [Supplementary Fig. S2c; t(6)=2.289, Student’s *t-test*].

**Figure 2.**
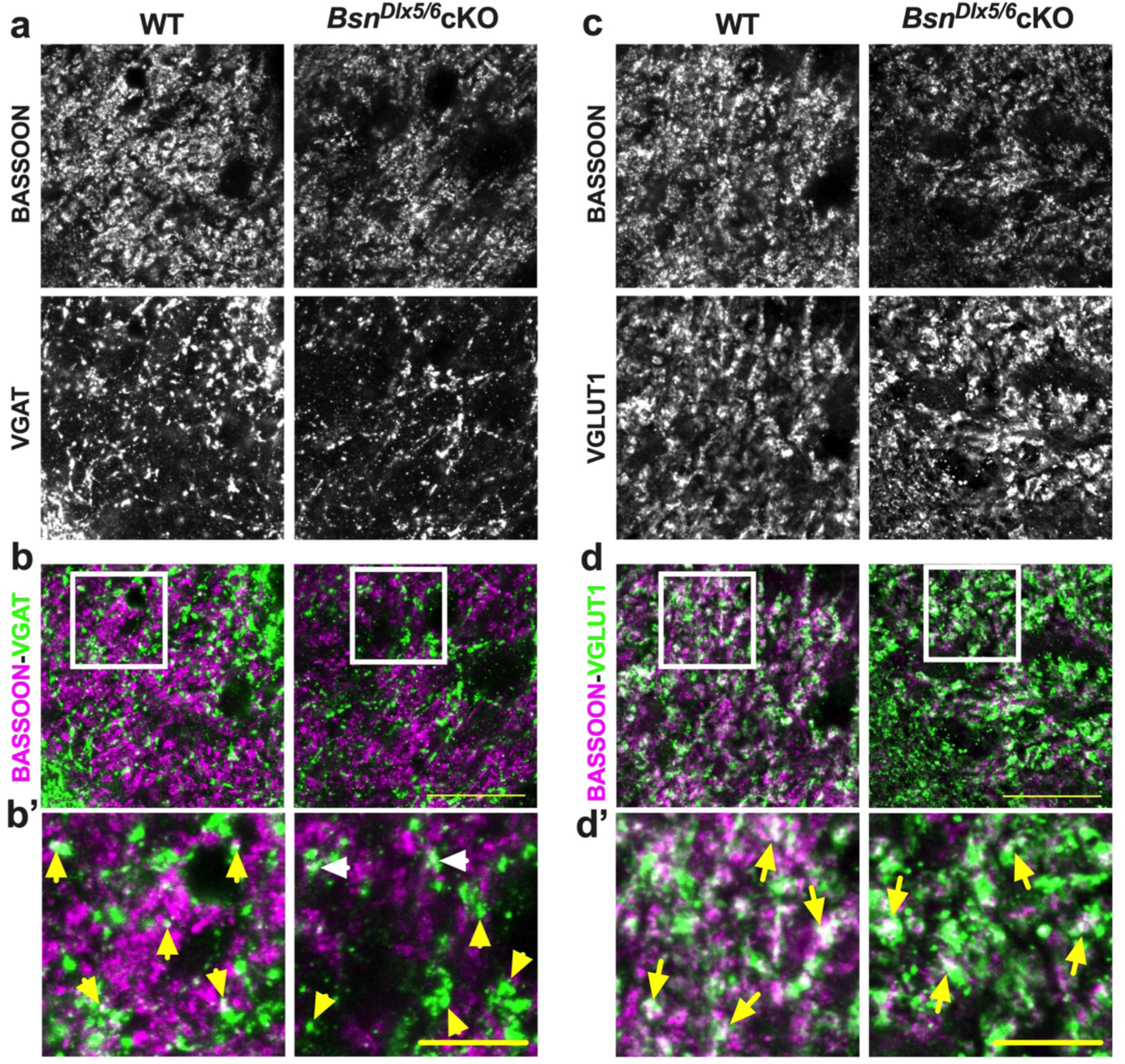
Lack of Bassoon at inhibitory presynapses of *Bsn^Dlx5/6^*cKO mice. **a)** Hippocampal sections (CA3 is shown) from WT and *Bsn^Dlx5/6^*cKO mice are stained with antibodies against Bassoon and VGAT. **b, b’)** Overlay and respective magnified images showing colocalization between Bassoon and VGAT in WT, but not in *Bsn^Dlx5/6^*cKO sections (indicated by yellow arrowheads). However, Bassoon expression is still present in some of the VGAT positive synapses of *Bsn^Dlx5/6^*cKO as indicated by white arrow heads. **c)** Another set of hippocampal CA3 sections from both the groups stained with Bassoon and VGLUT1 antibodies. **d-d’)** Overlay and zoom in images of WT and cKO sections documenting colocalization of Bassoon and VGLUT1 immunoreactivity in both the genotypes, as indicated by arrows. Scale bars in **b** and **d** are 25 µm and in **b’ and d’** is 10 µm.

Detailed densitometric analysis of the hippocampus revealed a reduced expression of VGAT and Synaptophysin in various layers of ventral dentate gyrus (DG) and CA3 subregions (Table 1, Supplementary Fig. S3).

**Table 1.** Immunohistochemical analysis in *Bsn^Dlx5/6^*cKO mice. Table 1: Immunohistochemical analysis of synaptic markers synaptophysin and VGAT in dorsal and ventral hippocampal areas in *Bsn^Dlx5/6^* cKO mice. Synaptophysin1 expression was found to be strongly reduced in ventral DG layers [(GCL: t(15)=4.991; IML: t(15)=7.493; OML: U=5]. Reduced expression of VGAT in various layers of ventral DG and CA3 subregions [(DG GCL: t(15)=3.537; CA3 SP: t(23)=2.392; CA3 SL: t(21)=2.094]. GCL=granule cell layer; IML=inner molecular layer; OML=outer molecular layer; SP=stratum pyramidale; SL=stratum lucidum; SO=stratum oriens; SR=stratum radiatum. N= 3 cKO and 3 WT mice and 2-6 sections per animal. All values are mean ± standard error of mean (SEM); *P ≤ 0.05, **P ≤0.01, ***P ≤0.001 ****P ≤0.0001, Unpaired Student’s *t*-test, Mann-Whitney *U*-test.

| Marker (mean integrated density normalized to WT levels) |  |  | Synaptophysin1 | VGAT |
| --- | --- | --- | --- | --- |
| Hippocampal area |  |  |  |  |
| **DG** | Dorsal | GCL | = | = |
|  |  | IML | ↓* | = |
|  |  | OML | = | = |
|  | Ventral | GCL | ↓*** | ↓** |
|  |  | IML | ↓**** | = |
|  |  | OML | ↓** | = |
| **CA3** | Dorsal | SP | = | = |
|  |  | SL | ↓p=0.058 | = |
|  |  | SO | = | = |
|  | Ventral | SP | ↓p=0.064 | ↓* |
|  |  | SL | = | ↓* |
|  |  | SO | = | = |
| **CA1** | Dorsal | SP | = | = |
|  |  | SR | = | = |
|  |  | SO | = | = |
|  | Ventral | SP | = | = |
|  |  | SR | = | = |
|  |  | SO | = | = |

### *Bsn^Dlx5/6^*cKO mice show disturbed GABAergic transmission and altered network activity

#### Disrupted inhibitory neurotransmission onto pyramidal cells in the ventral CA1 (vCA1 PCs)

We next tested for alterations in inhibitory and excitatory inputs onto vCA1 in acute hippocampal slice preparations of *Bsn^Dlx5/6^*cKO mice. The frequency of miniature inhibitory postsynaptic currents (mIPSCs) arriving at vCA1 pyramidal neurons was significantly lower in the *Bsn^Dlx5/6^*cKO slices than in controls [Fig. 3 a,c; inter-event interval of mIPSCs: t(8.2)=-2.59, Welch *t*-test]. No difference between the two genotypes was observed in the average amplitude of isolated mIPSCs [Fig. 3b; t(17)=-0.053]. Furthermore, conditional ablation of Bassoon had no effect on the kinetics of the recorded mIPSCs as the decay time constants of the events were similar in vCA1 PCs of both genotypes [Supplementary Fig. S4a; t(17)=-1.203]. These results suggest a reduction of inhibitory synaptic connections onto vCA1 pyramidal cells of the *Bsn^Dlx5/6^*cKO mice, while the postsynaptic compartment and the kinetics of these events remained unchanged. At the same time, we did not observe significant changes in the frequency of excitatory postsynaptic currents (mEPSCs) (Supplementary Fig. S4b, d; inter-event interval of mEPSCs, U=227, Mann-Whitney *U*-test) and/or in mEPSC amplitude in *Bsn^Dlx5/6^*cKO animals compared to control animals [Supplementary Fig. S4c; t(32.5)=-1.406].

**Figure 3.**
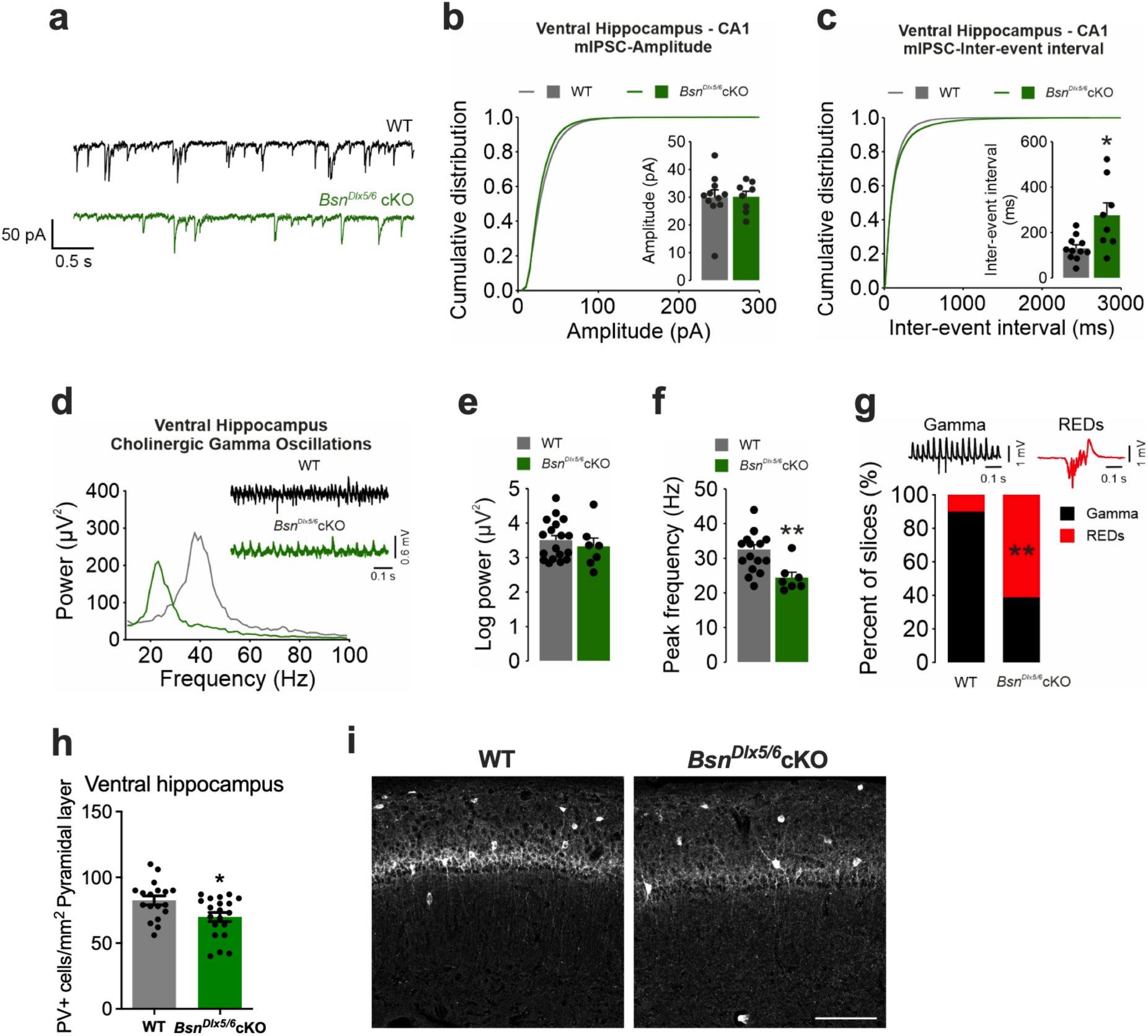
Disturbed inhibitory network configuration in the ventral hippocampus of *Bsn^Dlx5/6^*cKO mice. **a)** Representative traces of mIPSC recordings of vCA1 PCs from WT (black) and *Bsn^Dlx5/6^*cKO (green) animals. Membrane potential was held at -70 mV, mIPSCs were isolated by application of TTX (1 µM), CNQX (20 µM), and D-AP5 (50 µM), cells were recorded using high CsCl concentration based internal solution. **b)** Cumulative distribution of the amplitude of mIPSCs recorded in pyramidal cells of the ventral CA1. (Inset) Barplot is showing the mean ± SEM of the average mIPSC amplitude values of individual recorded cells (WT: 29.99 ± 2.63 pA; n=11 cells from N=3 mice; *Bsn^Dlx5/6^*cKO: 30.17 ± 1.94 pA; n=8 cells from N=3 mice). **c)** Cumulative distribution of the inter-event intervals of the recorded mIPSCs. (Inset) Mean ± SEM of the average mIPSC inter-event interval values of individual recorded cells (WT: 130.26 ± 15.83 ms, n=11 cells from N = 3 mice; *Bsn^Dlx5/6^*cKO: 276.37 ± 54.15 ms; n=8 cells from N=3 mice). Note that the significantly higher inter-event interval values of *Bsn^Dlx5/6^*cKO recordings indicate less functional inhibitory synaptic input to vCA1 PCs compared to WT (Welch *t* test, t(8.2)=-2.59, p=0.031). **d)** Representative power spectra of gamma oscillations from CA3 of WT and cKO mice showing a strong shift in the gamma peak frequency. Note the reduced number of gamma cycles in the cKO gamma trace (green) in comparison to WT gamma trace (black). **e-g)** Summary graphs showing no alteration in the gamma power (WT: N=5 mice, n=18 slices; cKO: N=5 mice, n=7 slices), a strong reduction in the gamma peak frequency (WT: N=5 mice, n=18 slices; cKO: N=5 mice, n=7 slices) and an increase in the number of recurrent epileptiform discharges (REDs) after Cch in the interneuron-specific cKO mice. **h-i)** Reduced number of PV interneurons in ventral hippocampus pyramidal layer of cKO mice (WT: n=18 sections from N=5 mice; *Bsn^Dlx5/6^*cKO: n=18-20 from N=5 mice). Cell numbers were counted manually using the Cell Counter plugin from Image-J. Scale bar is 200 µm. \**P* ≤ 0.05, \*\**P* ≤ 0.01, Fisheŕs exact test.

#### Reduced baseline transmission and altered short-term plasticity in Bsn^Dlx5/6^cKO mice

Next, we aimed to assess whether these changes are accompanied by altered transmission and plasticity at the ventral SC-CA1 synapse. Intriguingly, at this synapse, a lower excitability was evident when I-O curves were compared. Both presynaptic FV amplitudes (Supplementary Fig. S5a) and postsynaptic fEPSP slopes (Supplementary Fig. S5b) were significantly reduced (see Supplementary tables S1 and S2 for statistical details). Transmission rate (fEPSP slope/FV amplitude) per slice was also reduced indicating an overall reduced synaptic efficacy of this synapse (Supplementary Fig. S5c). Short-term plasticity was assessed by a paired-pulse (PP) protocol with intervals ranging from 10 ms to 500 ms at SC-CA1 synapse. We detected a significant increase in the PP ratios specifically at 10 ms and 250 ms intervals (Supplementary Fig. S5d; see Supplementary table S3 for statistical details). We further observed a mild delayed synaptic fatigue in *Bsn^Dlx5/6^*cKO mice when assessing the reduction in the fEPSP slopes during the 1st HFS used for LTP induction (Supplementary Fig. S5e; see Supplementary table S4 for statistical details). Last, we assessed long-term potentiation (LTP) using HFS and TBS protocols (Supplementary Fig. S5f and S5g, respectively) but found no change in the LTP when fEPSP slope values 30-40 min after LTP induction were compared [HFS: t(13)=-1.455, Student *t*-test; TBS: U=168, Mann–Whitney *U*-test]. These data indicate that evoked responses at the SC-CA1 synapses are reduced in *Bsn^Dlx5/6^*cKO mice.

#### Altered network activity in interneuron-specific Bassoon cKO mice

Based on these findings, we further analyzed the cholinergic gamma oscillations in the ventral hippocampus induced by application of the cholinergic agonist carbachol (Cch; 5 µM). Forty-five to sixty min after Cch perfusion stable gamma oscillations at low gamma range (Peak frequency: ∼30-40 Hz) were observed (Fig. 3d). While no change was evident in the power of oscillations at gamma range [20-80 Hz; Fig. 3d, e; t(23)=0.684, Student’s *t*-test], a strong reduction in the gamma peak frequency [Fig. 3d, f; t(23)=3.406] was noted in the CA3 region of *Bsn^Dlx5/6^*cKO mice. We further observed a strong increase in the number of slices that generate recurrent epileptiform discharges (REDs) in the *Bsn^Dlx5/6^*cKO mice (Fig. 3g; 11 out of 18 slices: i.e., 61%; Fisher’s exact test; see also [26]), while only a small fraction of WT slices (2 out of 20 slices: 10%) showed REDs, indicating an increased propensity to generate REDs in the interneuron-specific cKO mice.

#### Reduction in the number of Parvalbumin-containing interneurons in Bsn^Dlx5/6^cKO mice

We further investigated the density of PV interneurons in the hippocampal subregions of *Bsn^Dlx5/6^*cKO in comparison to WT mice. Our results indicate an overall reduction of PV positive cells in pyramidal layer of ventral hippocampus (Fig. 3h, i; U=100, Mann–Whitney *U*-test).

### *Bsn^Dlx5/6^*cKO mice show behavioral features reminiscent of ASD

We next aimed to determine the effects of these GABAergic synaptic deficits on behavioral parameters reminiscent of neurodevelopmental and neuropsychiatric disturbances. Home cage activity showed typical diurnal activity in both wildtype and *Bsn^Dlx5/6^cKO* mice with increased activity during the 12-hrs dark phase [Supplementary Fig. S6a-c; F(23,299)=30.24, two-way repeated measures ANOVA]. There was no genotype effect [F(1,13)=0.06] or genotype x daytime interaction [F(23,299)=0.66]. However, *Bsn^Dlx5/6^*cKO mice did not build complex nests comparable to those of WT mice and reached a lower quality score (Fig. 4p, U=0; Fig. 4q), indicating a disturbance of innate home cage behavior.

**Figure 4.**
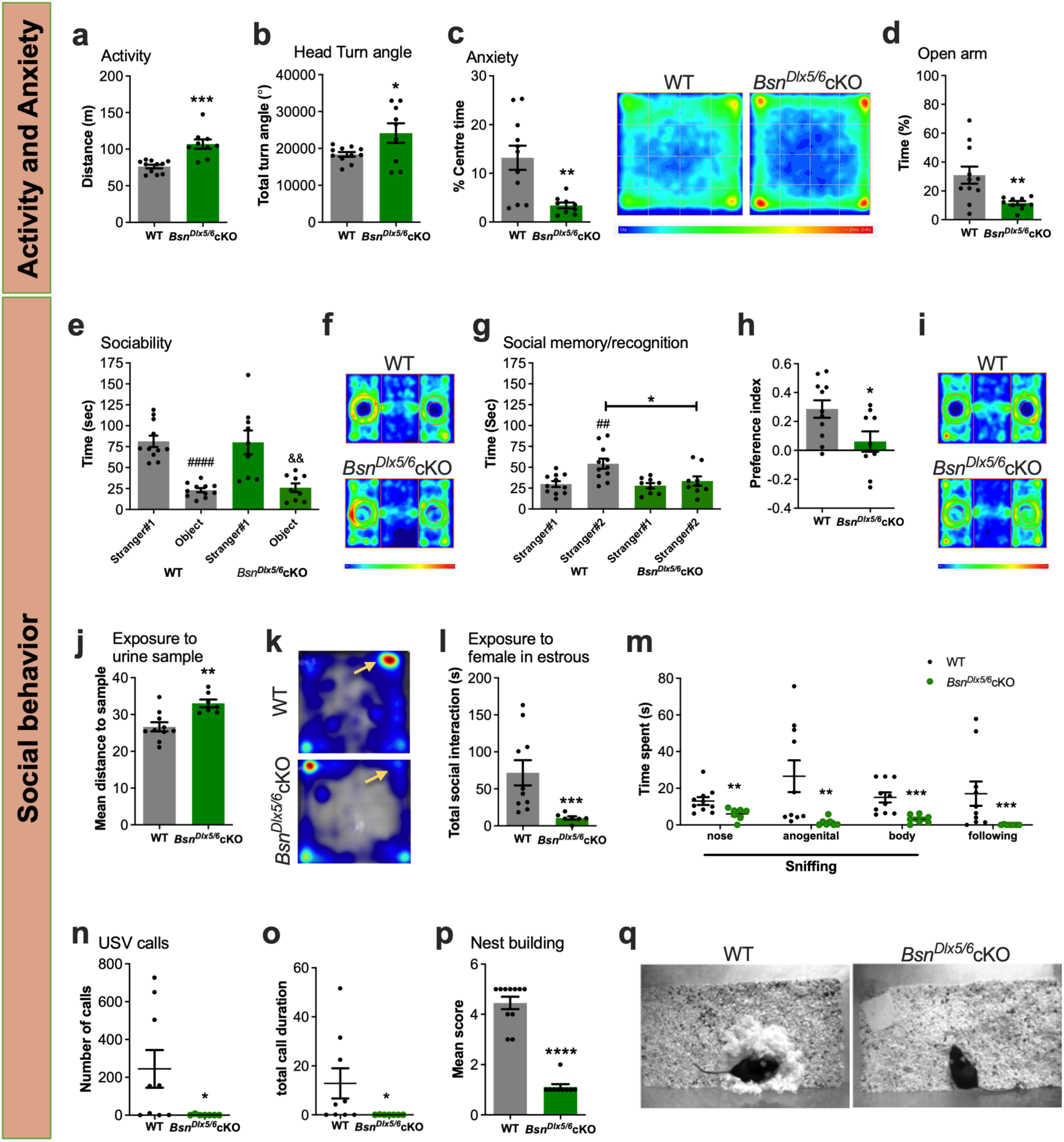
Emotional and social behavioral deficits reminiscent of ASD in *Bsn^Dlx5/6^*cKO mice. **a)** Increased activity (distance covered) of *Bsn^Dlx5/6^*cKO (N=9) mice compared to WT (N=11) in open field arena [50 cm (l) x 50 cm (w) x 35 cm (h). **b)** Increased total head turn angle in *Bsn^Dlx5/6^*cKO mice (N=9) compared to WT mice (N=11). **c)** Time spent by *Bsn^Dlx5/6^*cKO mice in center region of open field (25 cm x 25 cm), indicate anxiety like behavior compared to WT mice. Heat map images depicting the same during the session. Blue color indicates no activity (0 s) and red coloration indicates higher activity (2 min 24 s) pattern. **d)** During the elevated plus maze test, *Bsn^Dlx5/6^*cKO mice (N=9) spent less time in the open arm compared to WT mice (N=11). **e)** In a 3-chambered sociability test, both WT (N=11) and *Bsn^Dlx5/6^*cKO (N=9) mice prefer stranger #1 over a non-living object. **f)** Heat map images showing the mean group activity of WT and *Bsn^Dlx5/6^*cKO during the session. Blue color indicates no and red highest dwell time at a location. **g)** During social memory/recognition test, WT mice prefer new (stranger #2) over to familiar conspecifics (stranger #1). In contrast, *Bsn^Dlx5/6^*cKO mice failed to discriminate between novel and previously known mice. **h)** Heat map images showing the mean group dwell time of WT and *Bsn^Dlx5/6^*cKO during the session. **i)** WT (N=10) and *Bsn^Dlx5/6^*cKO (N=7) mice are exposed to urine samples of female mice in estrus in an open field with the dimensions 49 cm x 49 cm x 30 cm, placed in a sound-attenuated chamber; the mean distance to the sample is increased in *Bsn^Dlx5/6^*cKO mice. **j)** Heat maps showing the position of urine sample (arrows) and the color-coded dwell time of WT and cKO mice (blue low and red highest dwell time). **k)** When WT (N=10) and *Bsn^Dlx5/6^*cKO (N=7) mice were exposed to female mice in estrus, cKO mice spent much less time interacting with the females. **l)** Analyses of different types of social interactions including sniffing and following revealed significant reduction in all types for cKOs. **m, n)** Ultrasonic vocalizations recordings revealed a massive reduction in total number of calls and call duration in *Bsn^Dlx5/6^*cKO mice (N=7) compared to WT mice (N=9). **o)** Quantification of mean score (according to Deacon, 2006) revealed nest building deficits in *Bsn^Dlx5/6^*cKO mice (N=9) compared to WT (N=11). **p)** Example photographs showing nest building in WT and *Bsn^Dlx5/6^*cKO mice. All values are mean ± standard error of mean (SEM); \**P* ≤ 0.05, \*\**P* ≤ 0.01, \*\*\*\**P* ≤ 0.0001, two-way repeated measures ANOVA with Bonferroni post-test; Unpaired and paired Student’s *t* test, Mann-Whitney *U* test or χ^2^-test. “#” indicates comparison within the WT mice and “&” indicates comparison within the cKO mice.

#### Increased novelty-induced activity and altered exploratory behavior in Bsn^Dlx5/6^cKO mice

In an open field arena, *Bsn^Dlx5/6^*cKO mice displayed increased locomotory activity compared to WT mice (Fig. 4a. Distance: U=4, Mann–Whitney *U*-test; Mean speed: U=4.500, p=0.0002, data not shown) suggesting novelty-induced hyperactivity. This was accompanied by increased occurrence of rearing episodes in *Bsn^Dlx5/6^*cKO mice with shorter inter-event-intervals compared to WT mice [Supplementary Fig. S7c, d; t(18)=3.390; mean inter-event-interval (s) WT=10.74±1.13; *Bsn^Dlx5/6^*cKO=6.6±0.64). Alterations in observatory behavior were also noticed as *Bsn^Dlx5/6^*cKO mice showed increased absolute head turn angle [Fig. 4b; t(18)=2.312, Student’s *t*-test) and angular velocity [Supplementary Fig. S7a; t(18)=3.221) during the initial 5 minutes exploration in an open field arena. However, self-grooming events showed only a non-significant trend towards increase (Supplementary Fig. S7b; U=27, Mann–Whitney *U*-test). Together these observations describe novelty-induced hyperactivity with altered exploratory motor behavior in *Bsn^Dlx5/6^*cKO mice.

#### Bsn^Dlx5/6^cKO mice display elevated anxiety-like levels

The analysis of exploration patterns in the open field further revealed significant differences between the genotypes [center time (%): t(18)=3.470, Student’s *t*-test], with *Bsn^Dlx5/6^*cKO mice spending reduced time in the center indicating an increased anxiety-like behavior (Fig. 4c). Increased anxiety-like behavior was confirmed in the elevated plus maze, where *Bsn^Dlx5/6^*cKO mice spent significantly less time exploring the open arms (Fig. 4d; t(18)=2.899, Student’s *t*-test). However, there was no significant difference between genotypes in terms of proportion of open arm entries (%) (Supplementary Fig. S8a; t(18)=1.309). Furthermore, overall activity in elevated plus maze measured by total distance explored was again higher in *Bsn^Dlx5/6^*cKO compared to WT [Supplementary Fig. S8b; t(18)=3.880], in line with a novelty-induced hyperlocomotion of these mice. Furthermore, in the light-dark (L-D) test, *Bsn^Dlx5/6^*cKO mice spent less time in the illuminated compartment compared to WT mice (Supplementary Fig. S8c; U=9, Mann–Whitney *U*-test) and reduced number of transitions between compartments (Supplementary Fig. S8d; U=21.50). Again, *Bsn^Dlx5/6^*cKO also showed hyperlocomotion in this anxiety test (Supplementary Fig. S8e; t(13)=2.877).

#### Pronounced deficits in social behavioral exhibited by Bsn^Dlx5/6^cKO mice

Sociability and social recognition memory of male *Bsn^Dlx5/6^cKO mice* to same sex animals were assessed in a 3-chamber paradigm. Both genotypes displayed similar exploration behavior to the empty opaque cylinders during the habituation test [Supplementary Fig. S9a; WT: t(10)=1.033; *Bsn^Dlx5/6^*cKO: t(8)=0.2874, Student’s paired *t*-test]. During the sociability test, both genotypes displayed a preference for the social conspecific (Stranger#1) over the non-animated object [Fig. 4e, f; WT: t(10)=7.75; *Bsn^Dlx5/6^*cKO: t(8)=4.231]. During the social novelty test, however, *Bsn^Dlx5/6^*cKO mice in contrast to WT mice failed to prefer the novel stranger (Stranger#2) over the familiar one (Stranger#1) [Fig. 4g; WT: t(10)=4.068; *Bsn^Dlx5/6^*cKO: t(8)=1.208; Fig. 4h, i; WT vs *Bsn^Dlx5/6^*cKO at Stranger#2-Preference index: t(18)=2.434, Student’s *t*-test], indicating a social recognition or memory deficit in *Bsn^Dlx5/6^*cKO mice.

Next, social behavior of male *Bsn^Dlx5/6^*cKO mice towards the opposite sex was studied. The mean distance of male *Bsn^Dlx5/6^*cKO mice to an urine sample from a female in estrus was significantly increased compared to male WT mice [Fig. 4j, k; t(15)=3.70]. *Bsn^Dlx5/6^*cKO mice also spent significantly less time in the quarter of the open field with the sample zone (Supplementary Fig. S9b; U=10, Mann–Whitney *U*-test), although the latency to enter the sample zone was not affected by genotype (Supplementary Fig. S9c; U=23). Consistently, the total distance travelled during this exposure session was higher in *Bsn^Dlx5/6^*cKO mice [Supplementary Fig. S9d; t(15)=2.36]. Inspections of the acoustic data showed that there were only occasional ultrasonic vocalization calls not allowing for a detailed quantitative analysis. When analyzing the social interactions of male mice with female mouse in estrus we found a significant deficit in *Bsn^Dlx5/6^*cKO mice compared to WT mice (Fig. 4l; U=1). In line, a reduction was seen in the different types of social exploration (Fig. 4m), i.e., nose-nose sniffing (U=9), anogenital sniffing (U=5), body sniffing (U=2.5), as well as chasing behavior (U=4). Further, the proportion of sessions with ultrasonic vocalization was much lower in the sessions with *Bsn^Dlx5/6^*cKO mice than in those with WT mice (Supplementary Fig. S9e; χ^2^=6.35, Chi-square test) and the number of calls and the total call duration within sessions was reduced for those with *Bsn^Dlx5/6^*cKO mice (Fig. 4n, o; U=9 and U=10, Mann–Whitney *U*-test, respectively).

In summary, *Bsn^Dlx5/6^*cKO mice showed pronounced deficits in distinct social behaviors, both to same sex and opposite sex conspecifics, expressed in social exploration and memory and ultrasonic vocalization. We systematically controlled these behaviors as well as basic locomotor and anxiety-like behavior also in *Dlx5/6* driver mice, comparing *WT w/o Cre* (N=8) and *WT+Dlx5/6 Cre* (N=10) but did not find any overt phenotype (Supplementary Fig. S10), confirming that the observed behavioral changes in *Bsn^Dlx5/6^*cKO mice were indeed induced by the ablation of *Bsn*.

### Increased mitochondrial activity in *Bsn^Dlx5/6^*cKO mice

To investigate molecular mechanisms involved in the cellular and behavioral phenotype of *Bsn^Dlx5/6^*cKO mice, we performed proteomic analysis of hippocampal tissue. Comparison with WT samples revealed a total of 179 alterations in protein levels, 110 of which were up- and 69 down-regulated in the cKO (Fig. 5a, Supplementary table S5). Pathway enrichment analysis of the up-regulated proteins based on the MouseMine platform (open resource, [43]) showed that up-regulated proteins are mainly involved in the tricarboxylic acid (TCA) cycle and respiratory electron transport pathways (top two enriched pathways; Supplementary Fig. S11a), whereas no particular pathway was identified for the -down-regulated proteins. Gene ontology (GO) analysis identified association with mitochondria (Fig. 5b’, Supplementary table S6, S7) and metabolic/mitochondrial biological processes as most prominent associations (Fig. 5c’, Supplementary table S8, S9).

**Figure 5.**
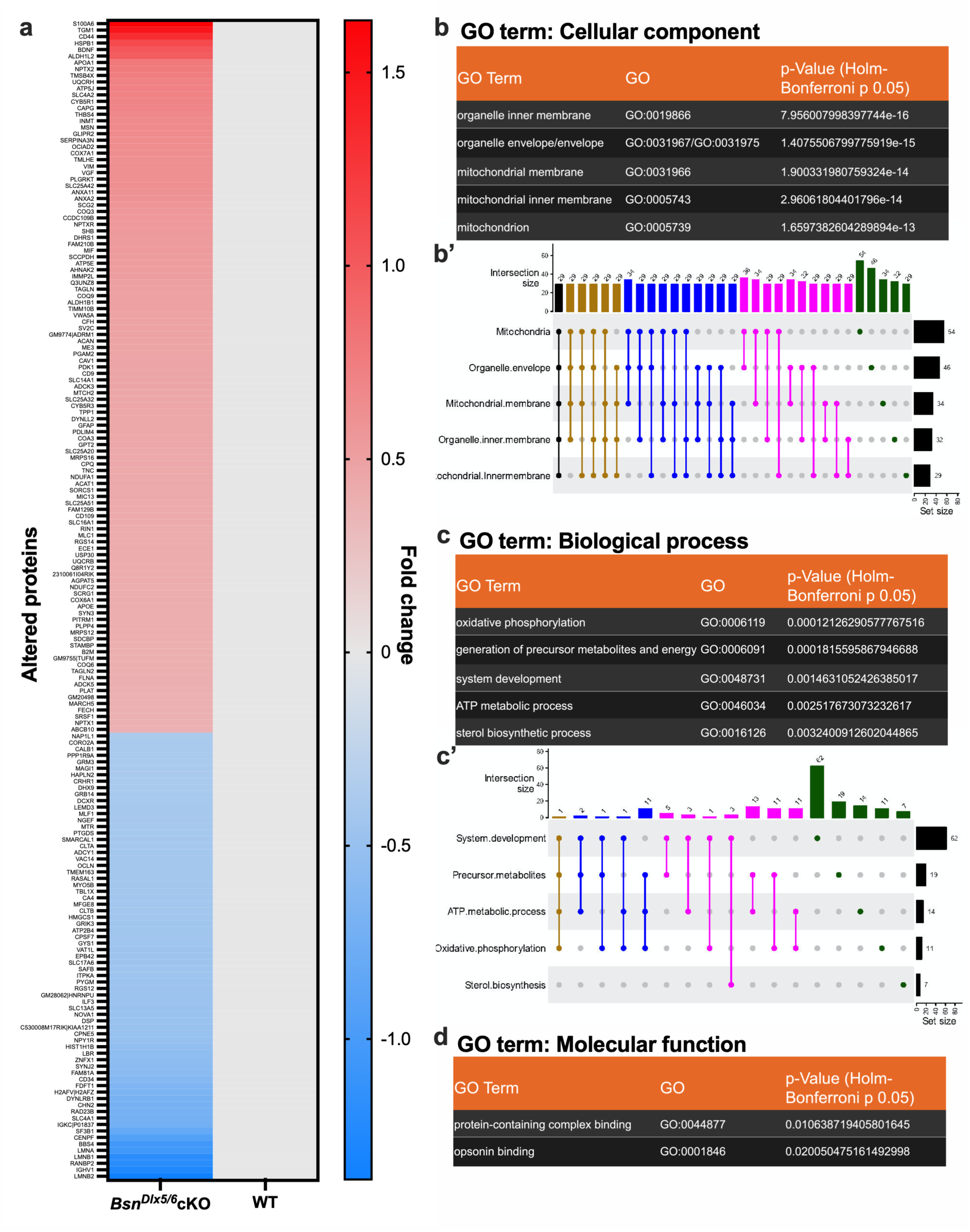
Altered levels of mitochondrial proteins in hippocampus of *Bsn^Dlx5/6^*cKO mice. **a)** Heat-map showing differentially regulated proteins in *Bsn^Dlx5/6^*cKO mice (N=4) normalized to WT (N=4), revealing changes in majority of mitochondrial proteins. The color code is a fold change (log2) values that indicates an up (in green) or down (red) regulation of different proteins. **b, c, d)** *Bsn^Dlx5/6^*cKO mice were compared to WT mice in GO cellular component, GO biological processes and GO molecular functions analyses, respectively. GO terms are mentioned in the order of decreasing enrichment p values (Holm-Bonferroni correction method) and the top 5 GO terms are mentioned here. **b’, c’)** UpSet plots indicate intersections or overlapping proteins among different enrichment terms within GO cellular components and GO biological processes, respectively. Set size on bottom-right side provides the information about total number of proteins in a specific enrichment term. Intersection size on y-axis indicates number of proteins that are overlapped or common among different enrichment terms. If there are ‘n’ number of proteins that are overlapped among all the top 5 GO terms, are indicated by a black-colored line, with highlighted black nodes in all the 5 respective GO terms. Gold/mustard-colored line represents number of proteins that are common only in 4 terms and the respective term is highlighted with same colored node. Blue-colored lines/nodes indicate proteins common among 3 terms, purple-color is for proteins common among 2 terms and green color nodes again represent the total proteins belongs to respective enrichment term.

GO analyses indicate a strong regulation of mitochondrial proteins (Fig. 6a) and related processes such as oxidative phosphorylation, respiration and ATP metabolism in *Bsn^Dlx5/6^*cKO. Targeted analysis furthermore identified seven ASD-(https://www.sfari.org/) and four schizophrenia-(SZ-) (http://szdb.org) associated candidates (Supplementary figures S11b,c), of which IMMP2L and TMLHE represent mitochondrial proteins.

**Figure 6.**
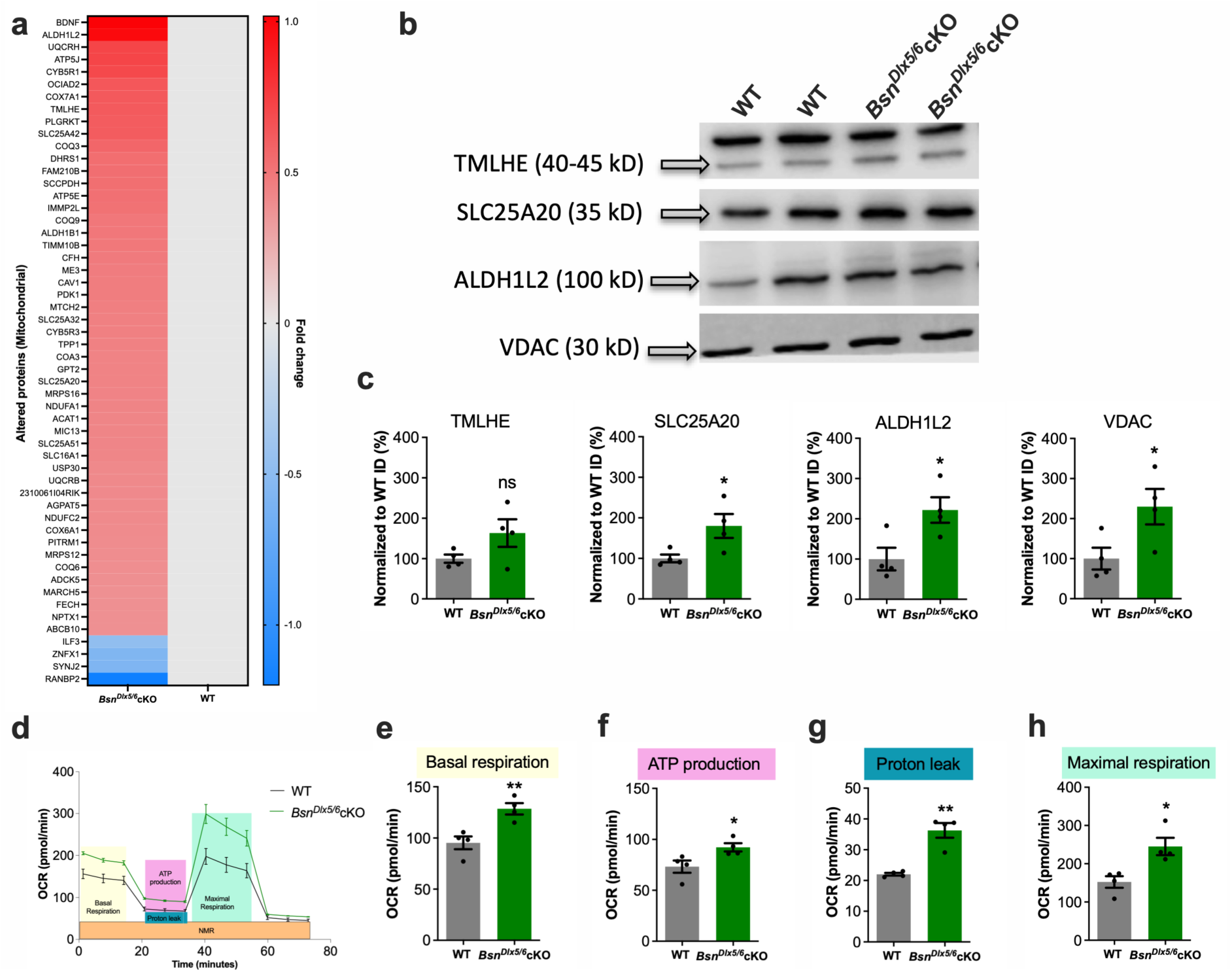
Upregulation of mitochondrial proteins and increased mitochondrial activity in *Bsn^Dlx5/6^* cKO mice. **a)** Heat-map displaying altered mitochondrial proteins in *Bsn^Dlx5/6^*cKO mice compared to WT. **b)** Representative Western blot images and **c)** quantifications for selected mitochondrial candidate proteins (TMLHE, SLC25A20, ALDH1L2) altered in proteomics screen and the mitochondrial marker VDAC. **d)** Mitochondrial respiration and key parameters were analyzed using Seahorse mito stress test on hippocampal cultures from WT and *Bsn^Dlx5/6^*cKO mice. Increase in overall mitochondrial kinetics measured as oxygen consumption rate (OCR) in *Bsn^Dlx5/6^*cKO mice. **e-h)** Increased mitochondrial activity resulting in basal respiration, ATP production, maximal respiration, and proton leak, suggesting a hyperactive mitochondrion in cKO mice. All values are mean ± SEM; \**P* ≤ 0.05. **P ≤ 0.01, Student’s *t*-test, Mann-Whitney *U*-test.

We confirmed altered expression of candidates that have been associated with ASD and are involved in mitochondrial metabolism including TMLHE, SLC25A20, ALDH1L2 as well as VDAC, a non-selective ion channel of the mitochondrial outer membrane, by immunoblot analysis (Fig. 6b). The tested proteins were found significantly up-regulated in *Bsn^Dlx5/6^*cKO mice, except for TMLHE, which did not reach statistical significance (Fig. 6b) [TMLHE: t(6)=1.770; SLC25A20: t(6)=2.585; ALDH1L2: t(6)=2.876; VDAC: t(6)=2.500, Student’s *t* test].

### Mitochondrial activity is increased upon Bassoon deletion

To examine related metabolic functions in *Bsn^Dlx5/6^*cKO mice, we quantified the oxygen consumption rate (OCR) in hippocampal primary cultures utilizing Seahorse technology. Neurons from cKO mice showed a significant increase in overall mitochondrial kinetics profile (Fig. 6d). Compared to WT neurons, basal and maximal respiration rates, along with ATP production-dependent respiration and proton leakage, were significantly increased in *Bsn^Dlx5/6^*cKO neurons [Fig. 6e-h; basal respiration: t(6)=3.961; ATP production: t(6)=2.626; maximal respiration: t(6)=3.383, Student’s *t* test; Proton leak: U=0, Mann–Whitney *U* test]. By contrast, no significant difference could be detected in non-mitochondrial respiration between genotypes [data not shown; t(6)=2.354, p=0.0567, Student’s *t*-test]. These data suggest an enhanced mitochondrial activity in *Bsn^Dlx5/6^*cKO primary neurons.

## Discussion

In this study, we demonstrate that Bassoon is critically important for the function of hippocampal GABAergic synapses and that a deficit in Bassoon in a subpopulation of such synapses in mice results in altered network configuration, metabolic imbalance and behavioral disturbances reminiscent of ASD.

The hippocampus has been proposed as an ideal structure to study in mice the synaptic and network processes underlying ASD-related behaviors and the role of specific genes involved [49]. *Bsn^Dlx5/6^*cKO mice display ASD core features (see Fig 4, Supplementary Fig. S7, S9) as well as co-occurring phenotypes of increased anxiety-like behaviors (Fig 4, Supplementary Fig. S8), adding to recently reported sensory learning deficits [50], decreased sensorimotor gating [51] and seizures (Fig 3g, [26]). Although additional brain regions certainly may be involved in the behavioral phenotype of *Bsn^Dlx5/6^*cKO mice, our hippocampus-directed analysis thus allowed us to examine relevant physiological correlates of their core ASD-related symptoms.

As a first step, we determined the inhibitory synaptic phenotype induced by Bassoon ablation. Bassoon has previously been implicated in the assembly of presynapses during development [4, 7]. However, we found that the ablation of *Bsn* in GABAergic interneurons does not evoke a loss of GABA synapses *in vitro* or in most layers of the hippocampus *in vivo*, suggesting that Bassoon may not be critical for the development and/or maintenance of inhibitory synapses *per se*. However, a selective loss of synaptic markers in the DG granule cell layer and CA3 pyramidal cell layer of *Bsn^Dlx5/6^*cKO mice indicates subfield-specific deficits. Moreover, reduced synaptic release from Bassoon-deficient inhibitory synapses in primary culture and reduced mIPSC frequency onto CA1 hippocampal pyramidal neurons as well as increased paired-pulse ratio could be observed strongly indicating a disturbance of GABA release.

Other animal studies have demonstrated the importance in the emergence of ASD-related behavioral deficits of inhibitory synapse-related genes, including the selective cell adhesion molecule neuroligin-3 [52, 53] and the intracellular signaling factors *ARHGEF9* [54] and Met [55] acting via gephyrin. Among different GABAergic interneuron populations PV interneurons and their activity have been most strongly associated with ASD [31, 32]. In fact, mutation of ASD candidate genes *CNTNAP2* [56, 57] and *Erbb4* [58] can impair differentiation of PV interneurons. In turn, disruption of PV interneuron function in mice can induce alterations of hippocampal network activity and behavioral features of ASD including altered social and exploratory behavior [59–61]. In line with these findings, we observed a reduction of PV cell counts in area CA1 of *Bsn^Dlx5/6^*cKO mice.

The subfield specific-changes of cellular and synaptic GABAergic markers in the hippocampus likely reflect a different hippocampal network organization of *Bsn^Dlx5/6^*cKO mice. Together with a general reduction in synaptic efficacy this argues for widespread changes of E/I balance and synaptic network configuration in the hippocampus of *Bsn^Dlx5/6^*cKO mice. Direct assessment of network activity patterns revealed frequency alterations in carbachol-induced gamma oscillations, likely related to altered PV interneuron function given their role in setting oscillatory pacing [62]. In contrast to *Bsn^ΔEx4/5^* KO mice, where synaptic plasticity deficits likely arise secondary to severe epileptic seizures [22], *Bsn^Dlx5/6^*cKO mice maintain largely intact long-term potentiation at CA3-CA1 synapses along with their epileptic phenotype [26]. However, *Bsn^Dlx5/6^*cKO mice exhibit enhanced short-term plasticity, which, despite the reduced baseline excitability, may contribute to their increased propensity for REDs and could be linked either directly or indirectly to their mild epileptic activity and PV interneurons dysfunction [63]. To begin to dissect the molecular consequences of interneuron-specific *Bsn* knockout, we investigated the hippocampal proteome, which surprisingly did not reveal any significant changes related to GABA synthesis, GABA receptors or the GABA shunt. Together, with altered GABAergic synaptic marker expression levels and physiological phenotype suggesting that *Bsn^Dlx5/6^*cKO mice display an altered hippocampal network organization rather than a general breakdown of GABAergic synapses. However, an increased expression of several mitochondrial proteins indicates a metabolic imbalance in *Bsn^Dlx5/6^*cKO mice, in agreement with observation in various ASD models [64] and possibly arising as consequence of the altered E/I balance [65, 66]. Amongst the upregulated ones we indeed found several metabolic pathway proteins that have been linked to ASD and/or schizophrenia (Supplementary Fig. S11b, c), such as TMLHE [67], the inner mitochondrial membrane protein IMMP2L [68] and voltage-gated anion channels (VDACs) [69]. We confirmed increased mitochondrial activity in *Bsn^Dlx5/6^*cKO cultures when measured for parameters like basal respiration, ATP production, and mitochondrial respiration. Furthermore, we observed an increase in proton leaks in *Bsn^Dlx5/6^*cKO neurons, in line with reports that have suggested that increased proton leakage is associated with some ASD models (reviewed by [64]). This altered proton leak may be related to the increase of voltage-gated anion channel (VDAC) protein seen in P2 fractions of the *Bsn^Dlx5/6^*cKO mutants.

Fast-spiking PV interneurons are considered extremely vulnerable to changes either in mitochondrial density or activity [70]; our data indicate that their malfunction may drive a vicious cycle of network activity change, metabolic imbalance and GABAergic disruption. Their high frequency of transmission makes these cells particularly dependent on P/Q-type Ca^2+^ channels for synaptic GABA release [71, 72]. Bassoon, via its second coil-coil domain is involved in the binding and selective positioning of the P/Q-type of Ca^2+^ channels at synaptic vesicle release sites, and therein indirectly (via Rim-binding proteins) interacts with the ASD candidates Rim1/2 [6]. They share this function with another ASD candidate Neurexin1a [73], a presynaptic ligand of Neuroligin-3 at interneuron synapses [74], which has moreover been identified as a putative interaction factor of Bassoon in a proximity labelling study [75].

A recent study examined the association of variants in *BSN* gene [12]. Of 29 patients that displayed epilepsy, five were diagnosed with autism or autistic-like behavior. The corresponding mutations related to a Pro207His mutation in the first coiled coil domain and various truncation mutations, covering a large range of position in *BSN* (at Thr535Profs*7, Thr1380Profs*19, Tyr2228* and Thr2720Alafs*38). Another recent study [13] has reported a protein truncated variant due to frameshift mutation in *BSN* (Arg1659Glnfs*23), in a child having ASD along with mild intellectual developmental disorder and epilepsy. We previously showed that in mice selective ablation of *Bsn* in excitatory forebrain neurons did not produce an ASD-like phenotype but rather increased baseline transmission at MPP-DG and SC-CA1 synapses, sharply contrasting our current observations in *Bsn^Dlx5/6^*cKO mice [27, 28]. It will be highly informative to determine how particular *BSN* mutations linked to ASD in humans may influence the function of GABAergic interneurons.

## Acknowledgements

We thank animal house employees from both LIN, IBIO and IPT for their wonderful service in taking care of the mice and technical assistant staff of LIN and IBIO. We thank Beat Lutz, Johannes Gutenberg-University Mainz, for advice on *Dlx5/6*-Cre mice.

## Contributions

Conceptualization, supervision of the study: A.A., E.D.G., O.S.

Cell biological experiments and analysis: C.M.-V.

Immunohistochemistry: A.A.

Metabolic analysis: N.T. and C.M.-V.

Design of mouse mutants: A.F.

Mouse genetics & breeding supervision: C.M.-V., A.A.

Electrophysiology experiments and analysis. G.C., E.M., S.M.

Evaluation of Proteomics, Bioinformatics: A.B., A.A., M.R., L.P.P.

Behavioral experiments and analysis. A.A., J.C.K., M.F., O.S.

All authors wrote, revised, or discussed the manuscript and agreed on this version.

## Availability of data and materials

All associated data from this manuscript are available from the corresponding authors upon reasonable request.

## Abbreviations

ADHD: Attention deficit hyperactivity disorder
ASD: Autism spectrum disorders
*BSN*: Human Bassoon gene
*Bsn*: Mouse Bassoon gene
Cch: Carbachol
cKO: Conditional knockout
DG: Dentate gyrus
GFAP: Glial fibrillary acidic protein
PV: Parvalbumin
SV: Synaptic vesicle
Syp1: Synaptophysin 1
Syt1Ab: Synaptotagmin-1 antibody
VGAT: Vesicular inhibitory amino acid transporter
VGLUT1: Vesicular glutamate transporter 1

## Supplementary Figures

**Supplementary Figure S1.**
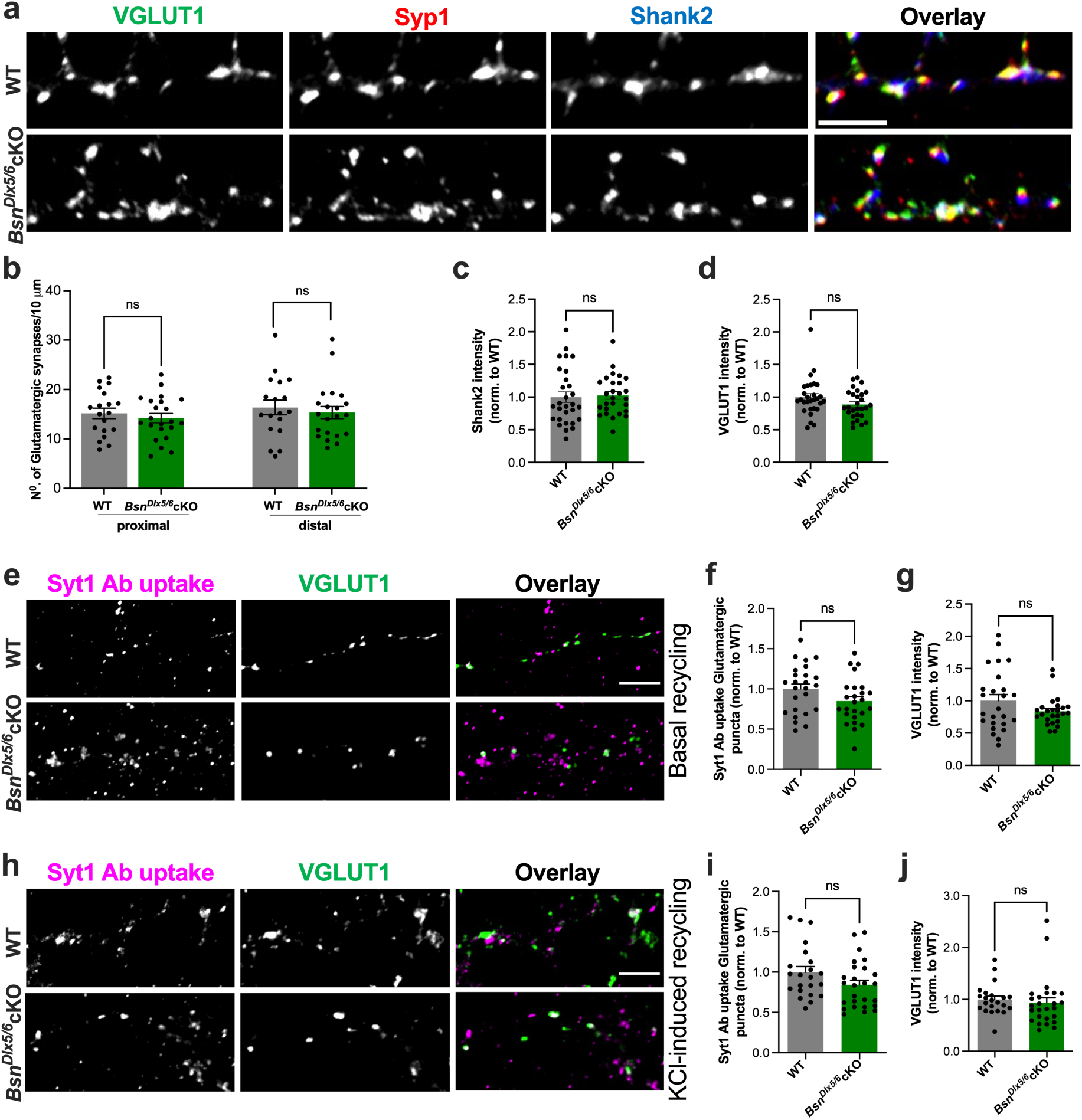
Synapse density and function of excitatory synapses are similar to wild types in *Bsn^Dlx5/6^*cKO hippocampal cultures. **a)** Representative images of 19-21 DIV cultured hippocampal neurons from WT and Bsn^Dlx5/6^cKO immunostained for VGLUT1, Syp1, and Shank2 to label excitatory synapses. **b)** Quantification of the number of excitatory synapses per 10 μm in the proximal and distal dendritic segments revealed no significant change between genotypes. **c,d)** Quantification of normalized IF intensity for Shank2 (**c**) and VGLUT1 (**d**) also revealed no significant changes. Data is obtained from 2 independent cultures per condition. All values are mean ± SEM. Statistical significance was assessed using Student’s *t* test as is depicted in graphs \**P* ≤ 0.05, \*\**P* ≤ 0.01, *** *P* ≤ 0.001. Scale bars, 5 μm. **e**) Representative images of Syt1Ab uptake (magenta) driven by endogenous network activity in hippocampal neurons 19-21 DIV from *WT* and *Bsn^Dlx5/6^cKO* mice. VGLUT1 (green) marks the excitatory synapses. **f)** Quantification of normalized immunofluorescence of Syt1Ab in excitatory synapses from experiments shown in. panel in e. **g)** Quantification of immunofluorescence intensity for Vglut1 showed no changes in the *Bsn^Dlx5/6^cKO* neurons. **h)** Representative images of Syt1Ab uptake (magenta) upon depolarization with 50mM KCl. Same cultures and staining conditions as in panel e. **i)** Quantification of normalized immunofluorescence of Syt1Ab uptake in excitatory synapses in experiments shown in panel h. **j)** Quantification of immunofluorescence intensity for VGLUT1 on experiments from panel h. Data are obtained from 3 independent culture preparations. All values are mean ± SEM. Statistical significance was assessed using Student’s *t-test, as* depicted in graphs \**P* ≤ 0.05, \*\**P* ≤ 0.01, *** *P* ≤ 0.001. Scale bars, 5 μm.

**Supplementary Figure S2.**
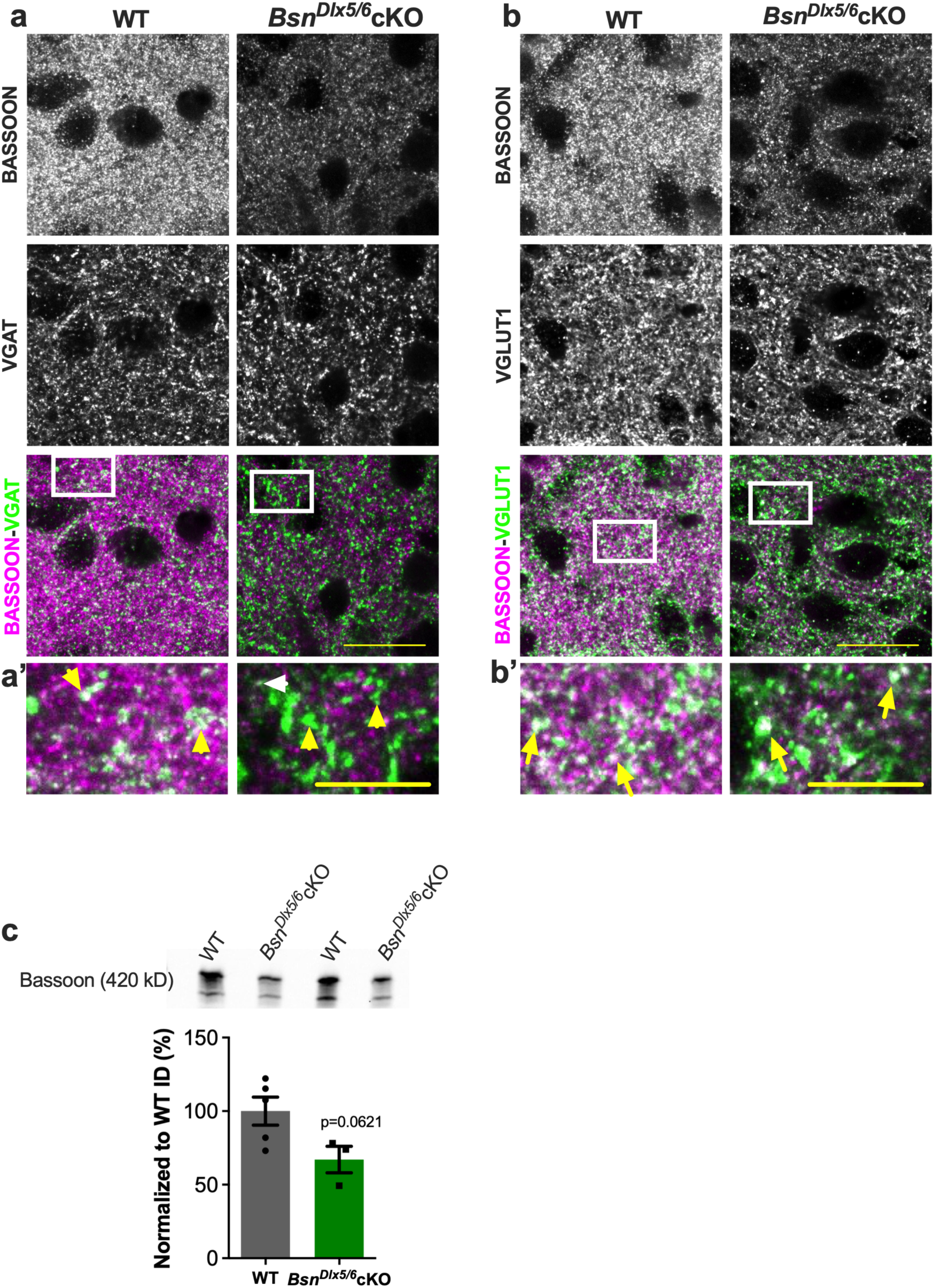
Characterization of Bassoon expression in brains of *Bsn^Dlx5/6^*cKO mice. **a)** Striatal sections from WT and *Bsn^Dlx5/6^*cKO mice stained with antibodies against Bassoon and VGAT. **a’)** Magnified overlay images showing colocalization of Bassoon and VGAT in WT but not in *Bsn^Dlx5/6^*cKO sections, as indicated by yellow arrow heads. Expression of Bassoon in some of the VGAT positive synapses of *Bsn^Dlx5/6^*cKO as indicated by white arrow heads. **b)** Striatal sections stained with antibodies against Bassoon and VGLUT1. **b’)** Magnified overlay images documenting colocalization between Bassoon and VGLUT1 immunoreactivities in both the genotypes (indicated by arrows). Scale bar in **a** and **b** are 25 µm and in **a’** and **b’** is 10 µm. **c)** Representative Western blot showing Bassoon and β-Tubulin (loading control) immunoreactivity analyzed using forebrain tissue from WT (N=5) and *Bsn^Dlx5/6^*cKO (N=3) mice and the respective quantification of the signal (WT: 100 ± 9.583%; *Bsn^Dlx5/6^*cKO: 67.12 ± 8.998%). All values are mean ± SEM; Student’s *t-*test.

**Supplementary Figure S3.**
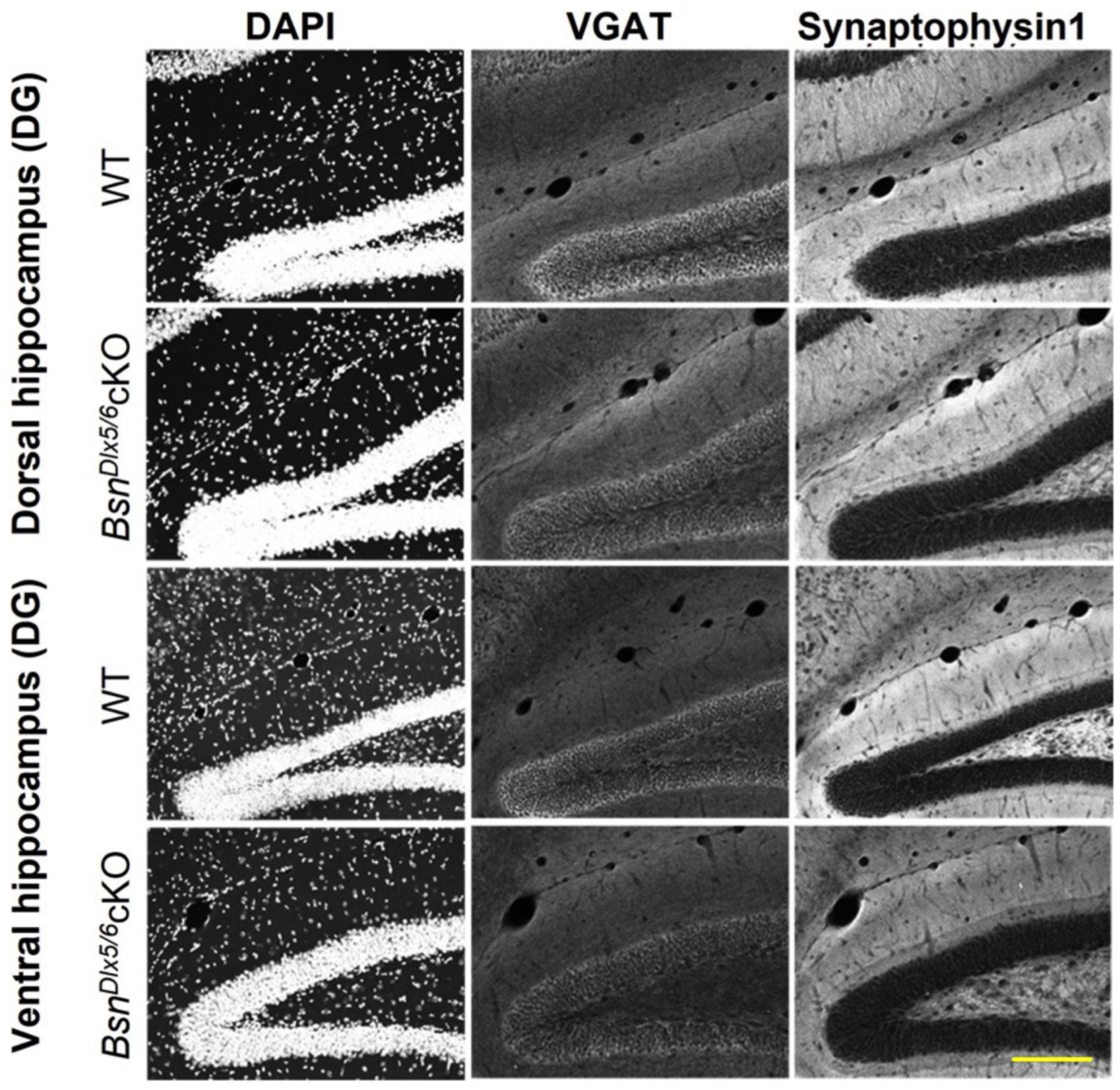
Immunohistochemical analysis of VGAT and Synaptophysin1 expression in *Bsn^Dlx5/6^*cKO brains. Representative images from WT and *Bsn^Dlx5/6^*cKO mice showing staining patterns of VGAT and synaptophysin1 in dorsal and ventral hippocampal sections (dentate gyrus, DG, is shown as an example). DAPI staining is used to define the cellular layers like the granule cell layer in DG and pyramidal layers in CA3 and CA1, to analyze the VGAT and Synaptophysin1 intensities. Further, Synaptophysin1 is used to differentiate the molecular layer sub-divisions in DG, *stratum lucidum* and *stratum radiatum* layers in CA3 and CA1, respectively, using free hand tool in Image-J. Scale bar is 200 µm.

**Supplementary Figure S4.**
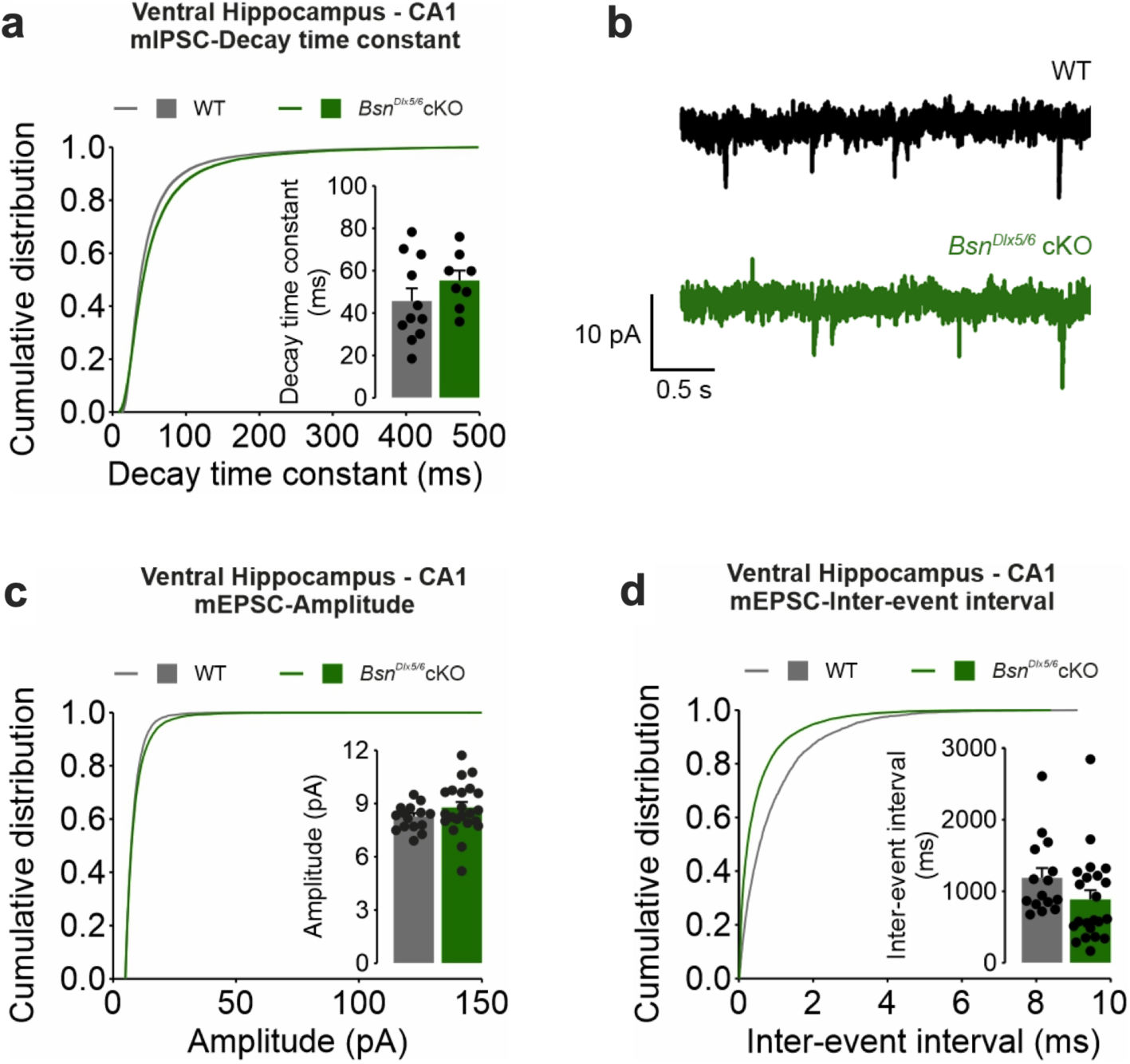
Unaltered mEPSC in the ventral CA1 pyramidal neurons of *Bsn^Dlx5/6^*cKO mice. **a)** Cumulative distribution of the decay time constant of the recorded mIPSCs. (Inset) mean ± SEM. of the average mIPSC decay time constant of individual recorded cells (WT: 45.68 ± 5.95 ms; n = 11 cells in N = 3 mice; *Bsn^Dlx5/6^*cKO: 55.37 ± 4.66 ms; n=8 cells from N=3 mice; t(17) = -1.203, p = 0.245). **b)** Representative traces of mEPSC recordings of vCA1 PCs from WT (black) and *Bsn^Dlx5/6^*cKO (green) animals. Membrane potential was held at -70 mV, mEPSCs were isolated by application of TTX (1 µM), and picrotoxin (100 µM), cells were recorded with low Cl-concentration internal solution. **c)** Cumulative distribution of the amplitude of mEPSCs recorded in pyramidal cells of the ventral CA1. (Inset) Barplot showing the mean ± SEM of the average mEPSC amplitude values of individual recorded cells (WT: 8.27 ± 0.18 pA; n=15 cells from N=7 animals; *Bsn^Dlx5/6^*cKO: 8.78 ± 0.31 pA; n=22 cells from N=6 animals; t(17) = -1.250, p = 0.22). **d)** Cumulative distribution of the inter-event intervals of the recorded mEPSCs. (Inset) Mean ± SEM of the average mEPSC inter-event interval values of individual recorded cells (WT: 1185.45 ± 138.24 ms; n=15 cells from N=7 animals; *Bsn^Dlx5/6^*cKO: 884.17 ± 130.59 ms; n=22 cells from N=6 animals; U = 103, p = 0.057).

**Supplementary Figure S5.**
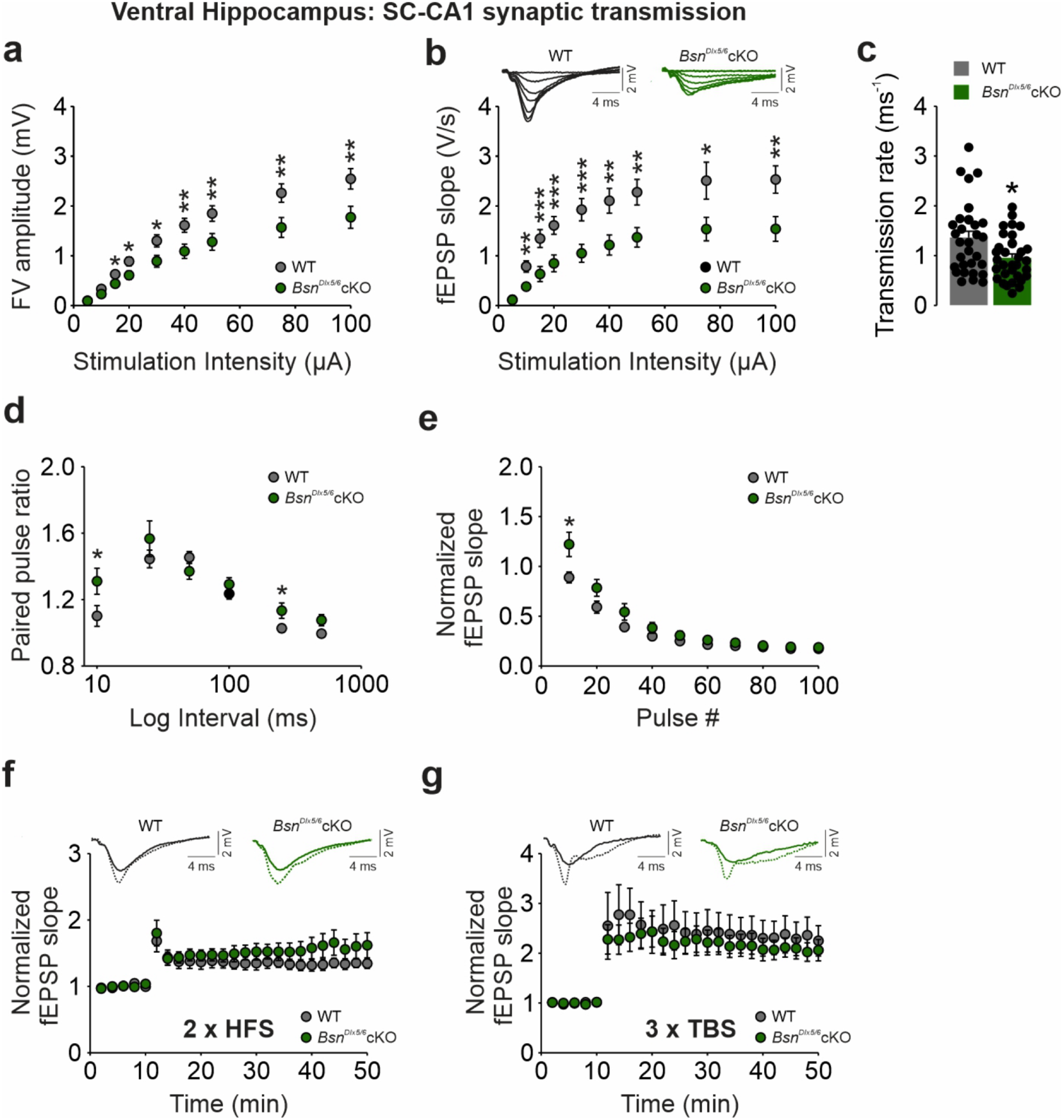
Reduced baseline transmission and enhanced short-term plasticity at the ventral SC-CA1 synapse of *Bsn^Dlx5/6^* cKO mice. **a)** Input-output (I-O) curves showing reduced FV responses (WT: N=11 mice, n=32 slices; cKO: N=11 mice, n=33 slices) and **b)** reduced fEPSP responses at the ventral SC-CA1 synapse of *Bsn^Dlx5/6^* cKO mice (WT: N=11 mice, n=32 slices; cKO: N=11 mice, n=34 slices). **c)** Summary graph indicating a reduced synaptic efficacy (reduced transmission rate per slice) at the ventral SC-CA1 synapse of cKO mice (WT: N=11 mice, n=32 slices; cKO: N=11 mice, n=33 slices). Summary graphs showing **d)** increased paired-pulse ratios (WT: N=11 mice, n=33 slices; cKO: N=11 mice, n=34 slices) at specific intervals (10 ms and 250 ms) accompanied by **e)** a mildly reduced synaptic fatigue during HFS (WT: N=4 mice, n=11 slices; cKO: N=4 mice, n=9 slices) in the ventral CA1 of *Bsn^Dlx5/6^* cKO mice. Summary graphs demonstrating no change in the **f)** HFS-induced LTP (WT: N=4 mice, n=8 slices; cKO: N=4 mice, n=7 slices) and **i)** TBS-induced LTP (WT: N=7 mice, n=17 slices; cKO: N=7 mice, n=20 slices. Data presented as mean ± SEM. **a, b:** Mann-Whitney *U* test or Student’s two-tailed test for each stimulation intensity; **d:** Mann-Whitney *U* test or Student’s two-tailed test for each stimulus interval; **e:** Student’s two-tailed test for each block of pulses; **c:** Mann-Whitney *U* test; **f:** Student’s two-tailed test; **g:** Mann-Whitney *U* test. \**P* ≤ 0.05, \*\**P* ≤ 0.01, *** *P* ≤ 0.001.

**Supplementary Figure S6.**
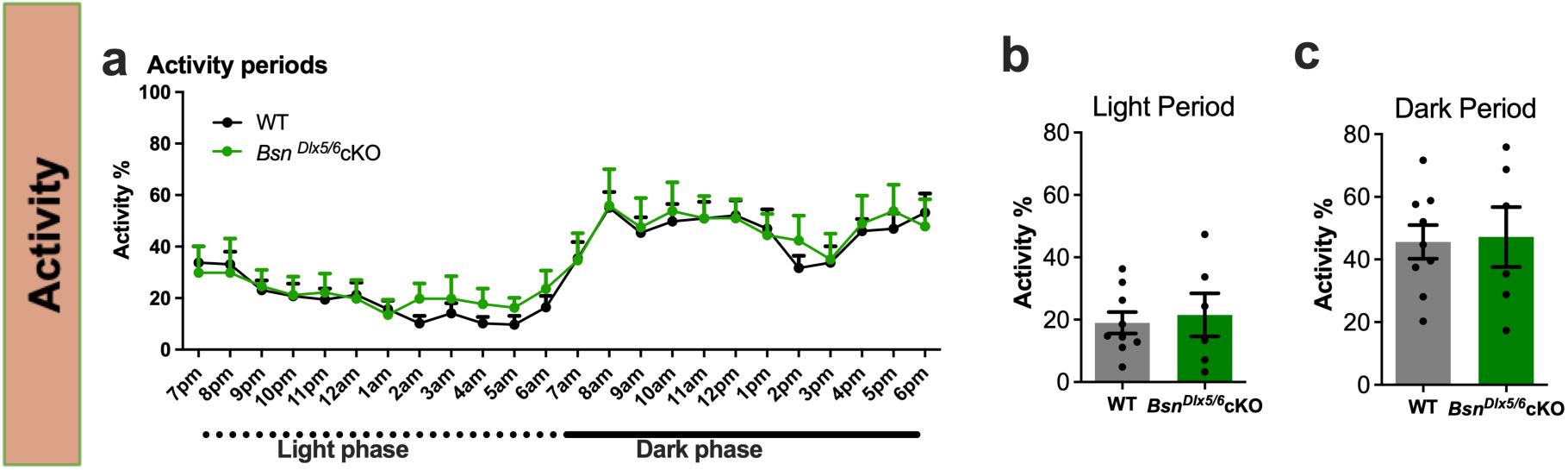
Normal home cage activity in *Bsn^Dlx5/6^*cKO mice. **a)** Home cage analysis of WT (N=9) and *Bsn^Dlx5/6^*cKO (N=6) display normal circadian activity and no genotype differences are observed. **b, c)** Both groups display low activity during light (Light phase activity (%) WT: 19.00 ± 3.452, cKO: 21.53 ± 6.908) and increased activity during dark period (dark phase activity (%) WT: 45.62 ± 5.389, cKO: 47.22 ± 9.577). All values are mean ± SEM; two-way repeated measures ANOVA with Bonferroni post-test, Student’s *t*-test.

**Supplementary Figure S7.**
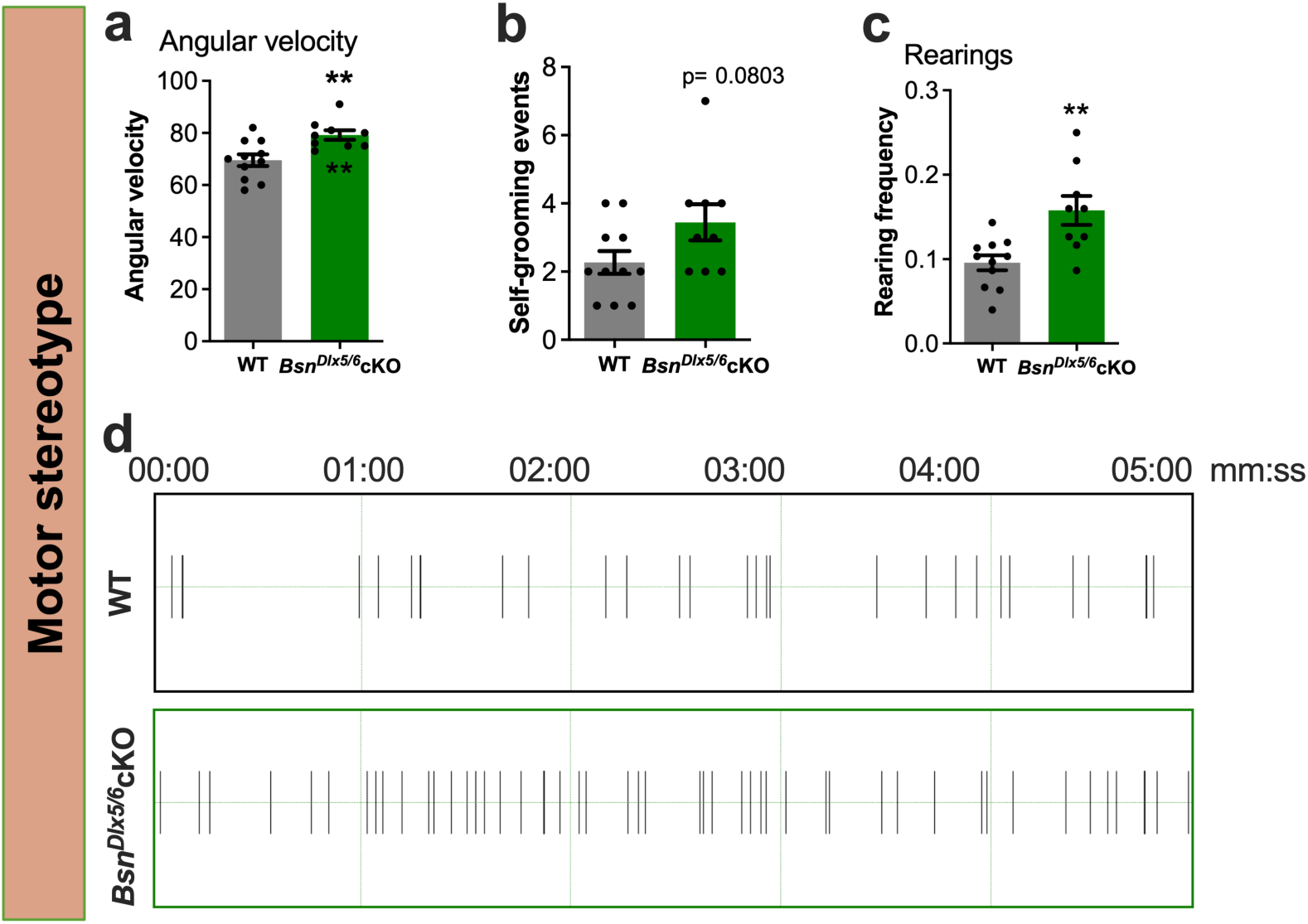
Analysis of motor patterns in *Bsn^Dlx5/6^*cKO mice. **a)** *Bsn^Dlx5/6^*cKO mice (N=9) display increased angular velocity compared to WT mice (N=11) and **b)** non-significant change in self-grooming events. **c)** Increased frequency of rearings (including rearings at the walls and free rearings) in open field arena in *Bsn^Dlx5/6^*cKO mice compared to WT mice. **d)** Representative picture displaying reduced inter-event-intervals of rearings in *Bsn^Dlx5/6^*cKO mice. All values are mean ± SEM; \**P* ≤ 0.05, **P ≤ 0.01, Student’s *t*-test, Mann-Whitney *U*-test.

**Supplementary Figure S8.**
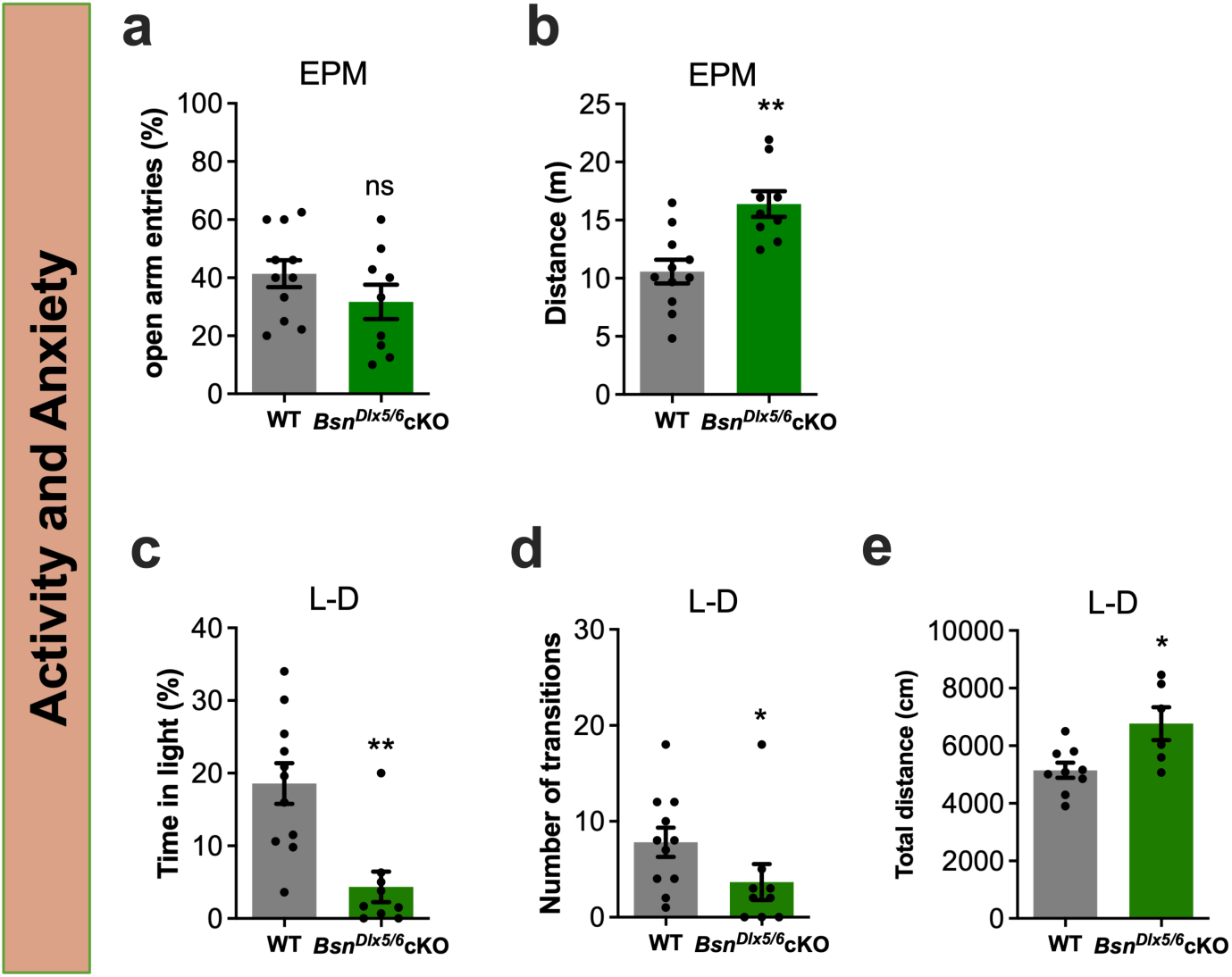
Assessment of activity and anxiety in *Bsn^Dlx5/6^*cKO mice. **a and b)** During the elevated plus maze (EPM) test, *Bsn^Dlx5/6^*cKO mice (N=9) display non-significant change in open arm entries (%), and display increased total activity compared to WT mice (N=11). **c and d)** This anxiety behavior is further confirmed in light-dark (L-D) test. The test chamber contains an illuminated chamber [19 cm (l) x 21 cm (w) x 20 cm (d)] and a dark chamber [17 cm (l) x 21 cm (w) x 20 cm (d)], connected by an opening (5 cm × 5 cm). *Bsn^Dlx5/6^*cKO mice spent reduced time in the illuminated compartment, and lesser number of transitions compared to WT mice. **e)** Further, *Bsn^Dlx5/6^*cKO mice (N=6) display increased total activity compared to WT mice (N=9), also during the L-D test. All values are mean ± SEM; \**P* ≤ 0.05, **P ≤ 0.01, Student’s *t*-test, Mann-Whitney *U*-test.

**Supplementary Figure S9.**
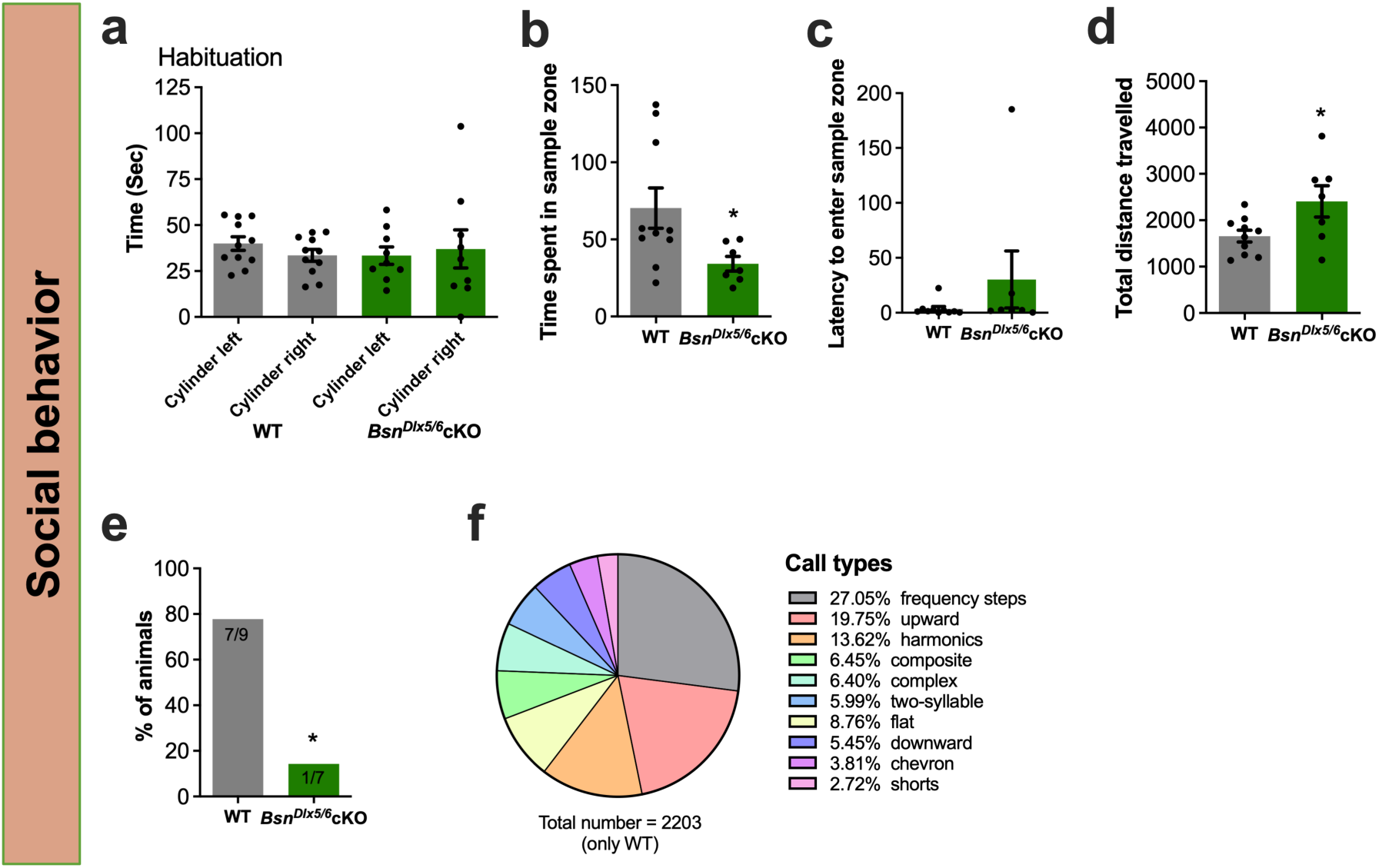
Social behavior of *Bsn^Dlx5/6^*cKO mice. **a)** During the habituation phase of a 3-chambered social test, both WT (N=11) and *Bsn^Dlx5/6^*cKO (N=9) mice explore the empty cylinders equally well without any discrimination. **b-d)** When exposed to urine of female mice in estrus *Bsn^Dlx5/6^*cKO (N=7) mice spent less time in the quarter of the sample compared to WT mice (N=9), no significantly different latency to enter this quarter was observed, and enhanced locomotor activity of *Bsn^Dlx5/6^*cKO mice was observed during this exposure session. **e)** During exposure to female mice in estrus, the proportion of sessions with ultrasonic vocalization (USV) was significantly reduced in *Bsn^Dlx5/6^*cKO mice (1 out of 7 mice elicit USV calls) compared to WT (7 out of 9 mice). **f)** Classification of the call types in the sessions with WT mice. The most prominent call types were frequency steps (27%), upward (20%) and harmonics (14%). All values are mean ± SEM; \**P* ≤ 0.05, Student’s *t* test, Mann-Whitney *U*-test, χ^2^-test.

**Supplementary Figure S10.**
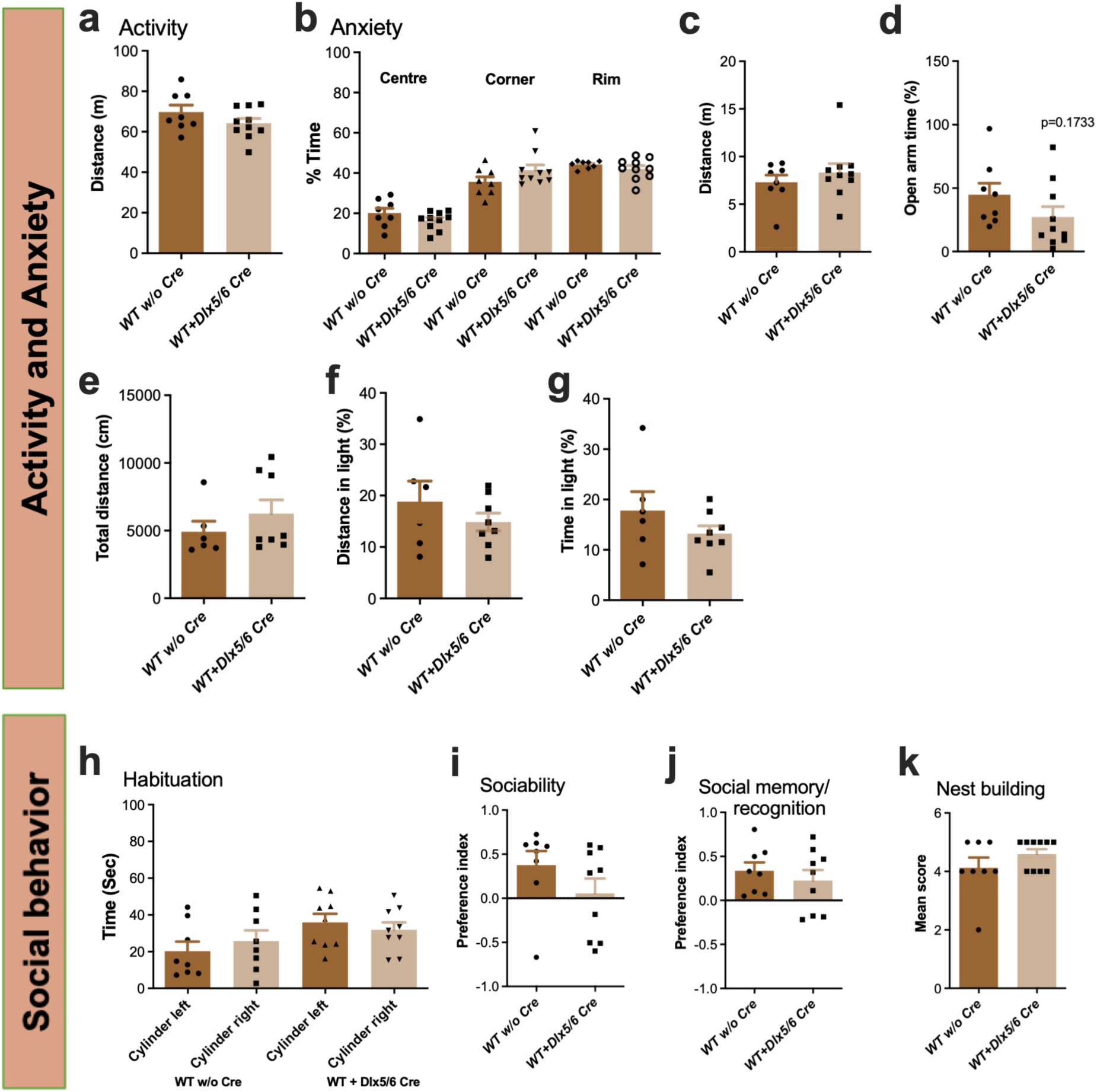
Expression of Dlx5/6-Cre transgene on *Bsn* wildtype background does not affect behavioral phenotype. **a)** In open field, both the groups, i.e. *Bsn^+/+^*(**WT w/o Cre**; N=8); and *Bsn^+/+^ X Tg(dlx5a-cre*) (**WT + dlxCre**): N=10), display similar exploratory behavior [t(16) = 1.375, p = 0.1882, Student’s *t*-test]. **b)** During time spent in different regions of open field by both the groups, significant effect of area, but neither genotype nor genotype x area interaction is observed in a 2-way repeated measures ANOVA with [Area: F(1.371, 21.94) = 57.56, p<0.0001; Genotype: F(1, 16) = 0.1540, p = 0.7000; Area x Genotype: F(2,32) = 2.117, p = 0.1369]. **c)** Total distance covered in EPM is not altered between the groups behavior [t(16) = 0.8207, p = 0.4239]. **d)** The open arm time (%), indicating an anxiety like behavior does not change between the groups [t(16) = 1.270, p = 0.2222]. **e)** Both the groups equally explored the chambers during the LD test (U = 15, p = 0.2824, Mann–Whitney *U*-test). **f, g)** Measure of activity in illuminated chamber is an indication for anxiety like behavior and both the groups covered equal distance [t(12) = 0.9963, p = 0.3388] and spent equal time [t(12) = 1.243, p = 0.2375], further indicating no change in anxiety like behavior due to Cre insertion. **h)** Social behavior was analyzed in a 3-chambered test. During habituation, both the groups spent equal time exploring the empty cylinders [*Bsn^+/+^*: t(7) = 0.8565, p = 0.4201; *Bsn^+/+^X Tg(dlx5a-cre*): t(8) = 0.4855, p = 0.6403 Student’s paired *t*-test]. **i, j)** Both control groups prefer the conspecific stranger (#1) during the sociability test over a non-animated object [t(15) = 1.368, p = 0.1914] and novel stranger (#2) over familiar stranger during the social memory or recognition test [t(15) = 0.7203, p = 0.4824], suggesting no obvious social phenotype among *Dlx5/6* control groups. **k)** Further in nest building-a measure of home cage social behavior, both the groups scored equally in building a quality nest (U = 29, p = 0.2878). All values are mean ± SEM; two-way repeated measures ANOVA, Unpaired and paired Student’s *t*-test and Mann–Whitney *U*-test.

**Supplementary Figure S11.**
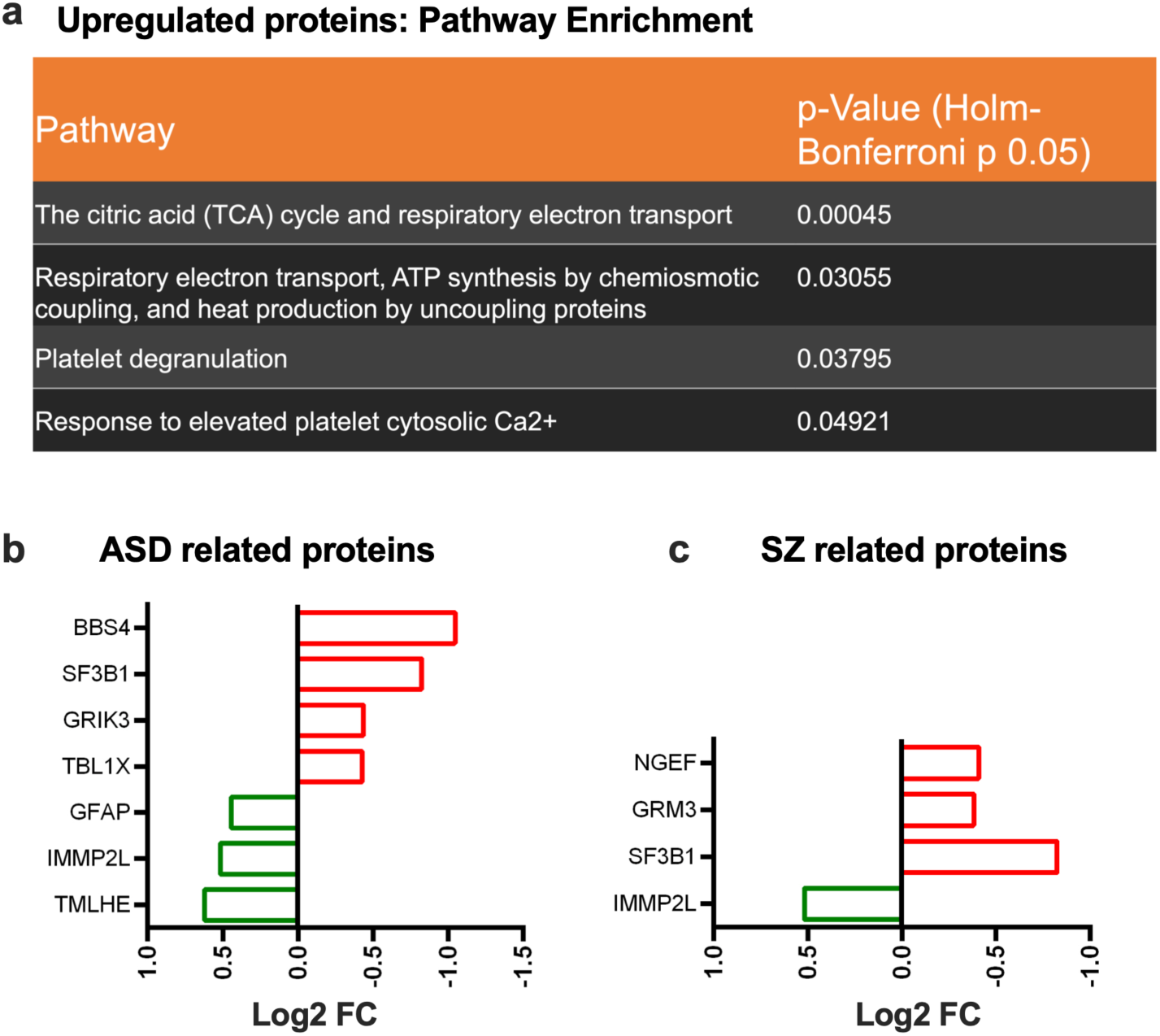
Pathway enrichment and neuropathological related proteins. **a)** The pathway enrichment analysis among upregulated proteins in *Bsn^Dlx5/6^*cKO mice, are associated with mitochondrial related functions. **b, c)** List of ASD (SFARI, https://www.sfari.org/) and SZ (http://www.szdb.org) associated proteins and their regulation (green color-upregulated and red color-down regulated) in *Bsn^Dlx5/6^*cKO mice.

