## Supplementary information for "Autism-like behavior induced by conditional ablation of the *Bassoon* gene in GABAergic interneurons"

### Supplementary materials and methods

#### Antibodies

Antibodies and antisera used in this study are listed in below table. Application details for Western blotting (WB), immunocytochemistry on primary neuronal cultures (ICC) and immunohistochemistry on brain slices (IHC) are indicated.

Table: List of antibodies used in the study

| Antibody | Host | Application | RRID | Manufacturer |
|---|---|---|---|---|
| Anti-ALDH1L2 | Rabbit | WB (1:1500) | RRID:AB_2878854 | Proteintech |
| Anti-Bassoon | Mouse | IHC (1:1000) | RRID:AB_10618753 | Enzo Life Sciences |
| | Rabbit | WB (1:1000) | RRID:AB_2313989 | Homemade |
| Anti-Gephyrin | Mouse | IHC (1:500) | RRID:AB_887717 | Synaptic Systems |
| Anti-Gephyrin | Rabbit | ICC (1:500) | RRID:AB_2619834 | Synaptic Systems |
| Anti-Bassoon | Guinea pig | ICC (1:500) | RRID:AB_2927388 | Synaptic Systems |
| Anti-PV | Mouse | IHC (1:1500) | RRID:AB_10000343 | Swant |
| Anti-SLC25A20 | Rabbit | WB (1:1500) | RRID:AB_10642001 | Proteintech |
| Anti-Synaptophysin1 | Rabbit | ICC (1:500)<br>IHC (1:1000) | RRID:AB_2864779 | Synaptic Systems |
| Anti-Synaptophysin1 | chicken | ICC (1:500) | RRID:AB_2622239 | Synaptic Systems |
| Anti-Synaptophysin1 | Mouse | ICC (1:1000) | RRID:AB_887824 | Synaptic Systems |
| Anti-Synaptotagmin1 | Rabbit | Live staining (1:70) | RRID:AB_887829 | Synaptic Systems |
| Anti-Shank2 | Guinea pig | ICC (1:500) | RRID:AB_2619861 | Synaptic Systems |
| Anti-TMLHE | Rabbit | WB (1:750) | RRID:AB_2205303 | Proteintech |
| Anti-Tubulin (Beta) | Mouse | WB (1:1000) | RRID:AB_477590 | Sigma-Aldrich |
| Anti-VDAC | Rabbit | WB (1:1000) | RRID:AB 10557420 | Cell Signalling |
| Anti-VGAT | Guinea pig | ICC (1:500)<br>IHC (1:500) | RRID:AB_887873 | Synaptic Systems |
| Anti-VGAT | Mouse | IHC (1:700) | RRID:AB_887872 | Synaptic Systems |
| Anti-VGLUT1 | Mouse | ICC (1:1000) | RRID:AB_262185 | Millipore |
| Anti-mouse Alexa 488 | Donkey | ICC (1:2000)<br>IHC (1:250) | RRID:AB_141607 | Invitrogen |

| | | | | |
|---|---|---|---|---|
| *Anti-mouse Cy3* | *Donkey* | *IHC (1:500)* | *RRID:AB_2315777* | *Jackson Immuno Research Labs* |
| *Anti-mouse Cy5* | *Donkey* | *IHC (1:250)* | *RRID:AB_2340820* | *Jackson Immuno Research Labs* |
| *Anti-rabbit 488* | *Donkey* | *IHC (1:250)* | *RRID:AB_141708* | *Invitrogen* |
| *Anti-rabbit Cy3* | *Donkey* | *ICC (1:2000)*<br>*IHC (1:250)* | *RRID:AB_2307443* | *Jackson Immuno Research Labs* |
| *Anti-guinea pig 488* | *Donkey* | *IHC (1:250)* | *RRID:AB_2340472* | *Jackson Immuno Research Labs* |
| *Anti-guinea pig Cy5* | *Donkey* | *ICC (1:1000)*<br>*IHC (1:500)* | *RRID:AB_2340462* | *Jackson Immuno Research Labs* |
| *Alexa Fluor 405 anti chicken* | *Goat* | *ICC (1:300)* | RRID:AB_2810980 | *Abcam* |
| *Anti-rabbit 680* | *Goat* | *WB (1:10000)* | *RRID:AB_10956166* | *Licor biosciences* |
| *Anti-rabbit 800* | *Donkey* | *WB (1:10000)* | *RRID:AB_2715510* | *Licor biosciences* |
| *Anti-guineapig HRP* | *Goat* | *WB (1:5000)* | ***RRID:***AB_2337402 | Jackson Immuno Research Labs |
| *Anti-rabbit HRP* | *Goat* | *WB (1:5000)* | *RRID:AB_228341* | *Thermo Fisher Scientific* |

#### Preparation of hippocampal primary neuronal cultures

For immunocytochemistry and live-cell assays, primary hippocampal cultures from newborn *Bsn*$^{Dlx5/6}$cKO and their WT littermates were prepared using the same protocol as described previously [1]. Briefly, after trypsin treatment and mechanical trituration, cells were plated in densities of 35000 cells per coverslip (18 mm diameter). One hour after plating, coverslips were transferred into dishes containing 60-70% confluent monolayer of astrocytes and Neurobasal A medium supplemented with B27, 11 mM sodium pyruvate, 4mM Glutamax, and antibiotics (100U/ml penicillin, 100ug/ml streptomycin). At 1 and 3 days, in vitro (DIV) Arac (Sigma Aldrich) was added to the cells (0,6 µM each time) to reach a final concentration of 1.2 µM. Neurons were maintained at 37 $^{0}$C in a humidified incubator containing 5% $CO_2$ until use.

#### Synaptotagmin1 luminal domain antibody uptake assay

Synaptotagmin1 luminal domain antibody (Syt1 Ab) uptake assays were performed as previously described [2]. Neurons were subsequently fixed with 4% PFA, permeabilized, and stained with antibodies against VGLUT1 and VGAT to discriminate excitatory and inhibitory boutons, respectively. To discriminate Bassoon-containing (Bsn+) from Bassoon-lacking (Bsn-) inhibitory presynaptic boutons in $Bsn^{Dlx5/6}$cKO hippocampal culture, co-staining with Bassoon antibody was applied.

#### Immunocytochemistry, image acquisition and quantification

Immunostainings of neuronal cultures were done as described in [2]. Briefly, primary hippocampal $Bsn^{Dlx5/6}$cKO and control neurons were fixed at 19-21 DIV. Then cells were permeabilized for 1 hour with blocking solution (10% fetal calf serum, 0.1% glycine, and 0,3% Triton X-100 in PBS) and incubated with primary antibodies (Table 1) overnight at $4^0$C, washed and incubated with secondary antibodies for 1h at RT (Table1) Afterwards, cells were dipped in water and mounted on glass slides with Mowiol.

Images for analysis of Syt1 Ab uptake in inhibitory presynaptic puncta were acquired using a Leica TCS SP8 STED 3X confocal microscope, equipped with a pulse White Laser (WLL) and a diode 405nm laser, using a 63X/1.40 NA Plan Apochromat oil objective. Image acquisition settings were: frame rate of 400 Hz with bidirectional scanning active, zoom of 2.5 and 1024x1024 resolution, achieving a final pixel size of 71.8 nm. For immunofluorescence (IF) quantification in each experiment, images from at least two different coverslips (10-12 cells each) were acquired and quantified to avoid effects given by experimental variance. The unspecific background was removed using threshold subtraction in ImageJ software. Afterwards, they were analyzed using the software ‘Openview’ (Tsuriel et al., 2006; updated version 4.0.1.0) kindly provided by N.E. Ziv. The function “Place Areas over Puncta” with a threshold of 500 was used to obtain the values for signal intensity. Images for analysis of Syt1 Ab uptake in excitatory presynaptic puncta and pre- and postsynaptic protein quantification were acquired on a Zeiss Axio Imager A2 microscope with Cool Snap EZ camera (Visitron Systems) controlled by VisiView (Visitron Systems GmbH) software. For quantifications, settings of camera or photomultiplier were applied identically to all coverslips quantified in one experiment. For IF quantification in each experiment, images from at least two different coverslips (7-10 cells each) were acquired and quantified to avoid effects given by experimental variance. The unspecific background was removed using threshold subtraction in ImageJ software. In all experiments, synaptic puncta were defined semi-automatically by setting rectangular regions of interest (ROI) with dimensions of about 0.8 X 0.8 µm around local intensity maxima in the channel with staining for synaptic marker Synaptophysin1, Shank2, Gephyrin, VGLUT1, or VGAT using OpenView software. Mean IF intensities were measured in synaptic ROIs in all corresponding channels using the same software and normalized to the mean IF intensities of the control group for each of the experiments.

#### Synapse density analysis

Densities of excitatory and inhibitory synapses were measured by counting the number of puncta in 20 µm along dendrites, proximal and distal (10-30 µm and 60- 80 µm away from the cell body, respectively). Dendritic segments co-stained with Synaptophysin1, VGLUT1, and Shank2 antibodies were used for the analysis of excitatory synapses. For assessing inhibitory synapse density, Synapthophysin1, Gephyrin, and VGAT antibodies were used. For synapse number analyses, ROI size was set to 5 x 20 µm. The individual presynaptic images were combined with their corresponding postsynaptic images to form a multi-channel image, saved in the required 16-bit format, and then examined for co-localization. To this end, we used the ComDet v.0.5.5 macro, which is available on GitHub (https://github.com/ekatrukha/ComDet). The analysis parameters were chosen as follows: maximum distance between co-localized spots: 7px, ROI shape: ovals. For each channel, the "include larger parameters" and "segment them" functions were enabled. The approximal particle size differs between excitatory and inhibitory proteins: For VGLUT1/Shank2, an approximal particle size of 7px was chosen, and for VGAT/Gephyrin, 9px. The intensity threshold was set to 12 SD in both cases.

#### Immunohistochemistry, image acquisition and analysis

Mice were perfused with PBS then with 4% PFA and postfixed overnight in same fixative. Brains were cryoprotected in sucrose solution and stored at −80 $^{0}$C until used for immunohistochemical staining [3]. Every 5th section (i.e., each 150 µm apart) from dorsal (−1.34 to −2.46 mm from Bregma) and ventral (−2.80 to −3.40 from Bregma) hippocampus was immunostained, employing different markers (Table 1). For quantification, confocal stacks of 0.2 μm (for staining intensities), 0.5 μm (for cell countings) with Z-step size (~2 and 5 μm Z-stack volume, respectively) were acquired with a Leica SP5 confocal microscope using 10x objective (HC PL Apo/0.40 CS/∞ 0.17/A) with 2x zoom factor and LCS software (Leica, Wetzlar, Germany) in different layers of sub-hippocampal regions including DG, CA3 and CA1. The analysed areas include: granule cell layer, inner molecular layer, outer molecular layer, stratum pyramidale, stratum lucidum, stratum oriens and stratum radiatum. Maximal projection images were obtained from the image stack using the Z-project function in Image-J software (https://imagej.net/ij/index.html).

#### Slice electrophysiology

##### Whole cell patch clamp recordings

Male $Bsn^{Dlx5/6}$cKO and WT mice (3-5 months old) were deeply anaesthetized with isoflurane before decapitating, and 300 μm horizontal hippocampal slices were prepared in ice-cold high sucrose artificial cerebrospinal fluid (85 mM NaCl, 75 mM sucrose, 2.5 mM KCl, 25 mM glucose, 1.25 mM NaH2PO4, 4 mM MgCl2, 0.5 mM CaCl2, and 24 mM NaHCO3, Leica VT1200S vibratome). After cutting, slices were incubated at 33°C for 30 min and then kept at room temperature until recordings were made. Recordings were performed in recirculated carbogen (95% O2, 5% CO2) saturated ACSF solution (125 mM NaCl, 3 mM KCl, 26 mM NaHCO3, 2.6 mM CaCl2, 1.3 mM MgCl2, 1.25 mM NaH2PO4, and 15 mM glucose) at room temperature. Fixed stage upright microscope (BX51WI; Olympus) equipped with infrared (900 nm) differential interference contrast optics (60x NA1W objective, Olympus) was used to visualize ventral hippocampal CA1 pyramidal neurons, and the cells were patched semi-randomly along the proximo-distal axis and the deep-superficial axis of the stratum pyramidale. Electrophysiological data were collected using ELC-03XS amplifier (NPI Electronic), InstruTECH LIH 8+8 16-bit Multi-Channel Data Acquisition System (HEKA) and by a custom written software in Igor Pro (WaveMetrics). Recording pipettes were pulled from borosilicate capillaries (inner diameter: 0.86 mm, outer diameter: 1.5 mm) using DMZ universal electrode puller (Zeitz-Instruments GmbH).

Pipettes were filled with a biocytin-containing intracellular solution for post hoc cell-type validation. For recording mEPSCs, low Cl- concentration internal solution was used (130.2 mM K-gluconate, 3.3 mM KCl, 1.8 mM NaCl, 1.7 mM MgCl2, 0.1 mM EGTA, 10 mM HEPES, 2 mM Mg-ATP, 0.4 mM Na2-GTP, 10 mM phosphocreatine, 8 mM biocytin; pH adjusted to 7.25 using KOH; 280–300 mOsm; pipette resistance: 3-5 MΩ), and 1 µM tetrodotoxin (Alomone labs), 100 µM picrotoxin (Tocris) and 0.12% DMSO (Sigma) was added to the ACSF solution. In mIPSC recordings, high Cl- concentration Cs+ based internal solution was used (133.5 mM CsCl, 1.8 mM NaCl, 1.7 mM MgCl2, 0.1 mM EGTA, 10 mM HEPES, 2 mM Mg-ATP, 0.4 mM Na2-GTP, 10 mM phosphocreatine, 8 mM biocytin; pH adjusted to 7.25 using CsOH; 280–300 mOsm; pipette resistance: 3-5 MΩ), and 1 µM tetrodotoxin (Alomone Labs), 20 µM CNQX (Tocris), 50 µM D-AP5 (Tocris) and 0.12% DMSO (Sigma) was added to the ACSF solution. The cells were held at -70 mV and to equilibrate the inner milieu of the cells, mEPSCs and mIPSCs were recorded 15 min and 20 min after the membrane break in, respectively. The sampling rate was 25 kHz, and every 10 s the series resistance of the recorded cell was monitored by applying a -5 mV voltage step. Data were discarded when the series resistance changed more than 20%.

After the experiments, slices were placed into fixative solution (4% paraformaldehyde in distilled water) overnight at 4°C. The specimens were then washed with PBS and blocked (1 h, 1% Triton X-100 in 1 M PBS, 10% NGS). Secondary antibodies (Alexa 488-conjugated streptavidin, in 1:500 concentration; Invitrogen, cat.no: S32354; RRID: AB_2315383) were applied for 2 h at room temperature in 0.5% Triton in PBS. Sections were mounted after several PBS washing steps by using Mowiol® 4-88 medium (Carl Roth Gmbh & Co. Kg). The recorded structures were visualized using an epi-fluorescent microscope (Axio Imager.A2, Zeiss). Amplitude, frequency, and time constant of mPSCs above 5 pA amplitude were analyzed using Matlab software (MathWorks) of the pyramidal cells with dendrites located in all strata of the ventral CA1.

#### Extracellular field recordings:

##### Slice preparation

Slice preparation and extracellular field recordings were performed similar to our previous studies [3–5]. Mice were decapitated and brains were extracted under deep isoflurane anaesthesia. Horizontal slices of ~400 µm thick brain slices were cut using an angled platform (12° in the fronto-occipital direction) in ice-cold, carbogenated (5% $CO_2$ / 95% $O_2$) artificial cerebrospinal fluid (aCSF) containing (in mM) 129 NaCl, 21 $NaHCO_3$, 3 KCl, 1.6 $CaCl_2$, 1.8 MgCl, 1.25 $NaH_2PO_4$ and 10 glucose (pH 7.4, ~300 mosmol / kg) with a vibrating microtome (Campden Instruments; Model 752) and quickly transferred to an interface chamber perfused with aCSF at 32 ± 0.5 °C (flow rate: 2.0 ± 0.2 ml /min). The ventral slices were cut until three-to-four slices containing ventral-to-mid hippocampus were obtained. Slices were incubated at least for an hour before recordings were started.

##### Evoked field potential recordings

Evoked synaptic responses from CA (Cornu Ammonis)3-to-CA1 synapse were obtained with a borosilicate glass electrode filled with aCSF (Resistance in aCSF: 1 MΩ) placed at the apical dendrites of area CA1 (Stratum Radiatum: SR) and a bipolar tungsten stimulation electrode (exposed tips: ~20 µm; tip separations of ~75 µm; electrode resistance in aCSF: ~0.1 MOhm) placed on the Schaffer collaterals (SC) at the proximal CA1 close to the CA2 subregion. A constant current stimulator was used to give square pulses with a stimulation duration of 100 µs. The responses were observed for a minimum of ten-to-twenty minutes to allow a stable recording (inter-stimulus interval of 30 sec). Nine pulses with a stimulation strength ranging from 5 to 100 µA were used to obtain an input-output (I-O) curve. Based on the I-O curve measurements, the stimulation intensity resulting in the maximal 40-50% evoked response was used for the rest of the experiment. To determine short-term plasticity, paired-pulse (PP) responses were recorded with intervals ranging from 10 to 500 ms. PP recordings were followed by a twenty-minute control recording (0.033 Hz) before commencing the LTP induction protocol. For LTP induction, either a high frequency simulation (HFS) protocol, a train of 100 pulses (100 Hz) repeated 2 times with 20 s interval or a theta-burst protocol (TBS: a train of 10 pulses (100 Hz) repeated 10 times with 200 ms interval) repeated 2 times with 10 s interval, was used. After LTP induction, responses were recorded for another 40 min (0.033 Hz). Extracellular field potential signals were pre-amplified using a custom-made amplifier and low-pass filtered at 3 kHz with a sampling rate of 5 kHz and stored on a computer hard disc for off-line analysis (Cambridge Electronic Design, Cambridge, UK).

Evoked field excitatory postsynaptic potentials (fEPSP) at SC-CA1 data were analyzed with MATLAB-based analysis tools (MathWorks, Natick, MA). For the analysis of fEPSP, the slope (V/s) between the 20 and 80% of the fEPSP amplitudes was measured. The presynaptic fiber volley (FV) was measured using the amplitude of the descending phase of the FV. To determine a measure for synaptic transmission efficacy, an average baseline transmission rate per slice was calculated by dividing each postsynaptic slope with the corresponding FV amplitude. The mean of these values was used to determine a transmission rate ($ms^{-1}$) per slice. PP ratios were calculated by dividing the slope of the second fEPSP to the first one. All LTP data, was normalized to the average of the baseline values obtained during 10 min before LTP induction.

#### Cholinergic gamma oscillations

Cholinergic gamma oscillations were induced via continuous bath perfusion of freshly-diluted carbachol (CCh, 5 µM). Before CCh perfusion, the temperature of interface chamber was set to 35°C to increase the likelihood of obtaining oscillations at gamma range (>30 Hz). After 45-to-60 min a glass electrode was placed at the CA3 pyramidal layer and three-to-five min recordings were obtained. Custom-made spike2 scripts were used to analyse gamma oscillations (Cambridge Electronic Design, Cambridge, UK). Power spectra were generated from two min artifact free data using Fast Fourier Transformation with a frequency resolution of 0.8192 Hz. Peak frequency (Hz) and Integrated power (20-80 Hz; $\mu V^2$) were calculated from the power spectra. Gamma recordings with peak powers lower than 50 $\mu V^2$ and peak frequencies lower than 20 Hz were discarded.

#### Behavioral experiments

##### Home cage activity (HCA) monitoring

One batch of male *Bsn*$^{Dlx5/6}$ cKO (N=9) and WT littermate mice (N=11) were habituated for two weeks to an inverted light cycle (lights on between 7PM to 7AM) and then subjected to a series of tests comprising HCA monitoring, open field, elevated plus maze, light-dark test, same sex interaction and social memory test and nest building.

HCA monitoring of the mice was performed for 4 consecutive days in their home cages using Home Cage Activity System (Coulbourn Instruments, Allentown PA). Activity was measured based on a threshold of movements between 100 ms (lower limit) and 500 ms (upper limit) for 15 s (raw values) and calculated the 5 min bins and further to one hour of activity periods [3]. Subsequent behavioral experiments in this batch were conducted between 10AM and 5PM, i.e. in the dark, active phase of the cycle.

##### Open field activity

Novelty-induced exploratory activity, anxiety-like behavior and motor stereotype behaviors were analysed in an open field arena, under low light (5 lux) conditions [3]. Using ANY-maze video tracking system (version 4.50, Stoelting Co, Wood Dale, IL, USA), the open field arena was divided into center, corners, and rims. The total distance moved, and the percentage of time spent in center region by each mouse, were measured and compared between the genotypes. Parameters like absolute head turn angle was measured and angular velocity was calculated by dividing absolute turn angle to the duration. Further, using BORIS (Behavioural Observation Research Interactive Software- a free and open-source software; [6]) the motor stereotype events like self-grooming, number and frequency of rearings (inter-event-interval) were analysed for initial 5 min.

##### Elevated plus maze (EPM)

Mice were further tested in EPM to measure anxiety levels. The experimental apparatus consists of total four arms with two closed arms [35 cm (l) x 5 cm (w) x 15 cm (h)] and two open arms [35 cm (l) x 5 cm (w)], which were placed at 40 cm above the ground surface. Experiment was performed under low light (5 Lux) for 5 minutes. Total distance, time spent and number of entries to the closed and open arms were tracked using ANY-maze Video tracking system (version 4.50, Stoelting Co, Wood Dale, IL, USA).

##### Light-dark (L-D) test

L-D test was conducted to measure anxiety-like behavior in mice in a test apparatus comprising an interconnected illuminated and dark chamber [3]. The activity (time spent), total distance (both chambers), and number of crossings in illuminated chamber were analyzed using the photo beams (TSE System, Bad Homburg, Germany) for 5 min as a measure of anxiety like behavior in mice.

##### Social interaction and memory

3-chambered social test is used to measure deficits in social interaction, social novelty or memory among the mice [7]. The test apparatus was made of a 60 cm (l) x 30 cm (w) x 40 cm (d) clear cage, separated by partitions for the mouse to explore the chambers. Empty cylinders made of wire with mesh-like holes [13 cm (h) x 4 cm (r)] are placed in lateral chambers during the test. During habituation, the test mouse was allowed to explore the empty cylinders for 5 min. During sociability or social interaction phase, a stranger mouse (C57BL/6J male) was placed in one of the cylinders and mouse-like animated object (made of Lego blocks) was placed in another cylinder. The test mouse was allowed to explore either a stranger mouse (Stranger#1) or an object for 5 min. The mesh-like holes of the wire cylinders allows both visual and olfactory communication. The final phase was social memory, where the object was replaced by a novel stranger mouse (Stranger#2), and the test mouse was allowed to explore familiar and novel stranger mice for 5 min. The activity in different phases, including exploration time exploring the cylinders, was measured by a video-camera and analyzed with the ANY-maze automatically and manually, and compared between the genotypes. The preference or discrimination index was measured using the formula "difference of exploration time between novel and familiar divided by total exploration time" and compared between the genotypes.

#### Nest building

Nest building activity was scored as described previously [8]. During this test each mouse in its home cage received a 2-3 gram weighted cotton nestlet. After one day, the scoring of 1-5 was given to each mouse based on the quality of the nest built by the mice representing a range from untouched nestlet to a complex nest with surrounding walls and a burrow.

#### Exposure to urine of females in estrus and females in estrus

Another group of male *Bsn*$^{Dlx5/6}$ cKO (N=7) and WT littermate mice (N=10) were tested under normal light cycle (lights on between 6AM to 6PM) regarding their behavioral responses during exposure to estrus female mice urine and to estrus females, including the ultrasonic vocalizations during the interaction with the female mice. For the exposures, an open field placed in a sound-attenuated chamber, was used. The behavior of the mice was recorded by a camera mounted on the ceiling of the chamber. Analyses of the behavior were performed automatically using a tracking software (Ethovision XL 11; Noldus, Wageningen, NL) and manually using an event recorder (Solomon Coder, https://solomon-coder.software.informer.com). Ultrasonic vocalization was monitored by an UltraSoundGate Condenser Microphone CM 16 (Avisoft, Berlin, Germany) mounted on one upper corner of the open field. The microphone was connected to an UltraSoundGate 116 USB audio device which was connected to a computer with Avisoft Recorder software. The acoustic data were recorded with a sampling rate of 250 kHz and were later analyzed with AviSoft SASLab Pro software (fast Fourier transformation with 512 FFT length, 100% frame, Hamming window and 75% time overlap). The calls were manually marked and categorized by an experienced user blinded to genotypes using the call classification published by Scattoni and colleagues [9]. For the exposures, female C57BL/6J mice at an age of 8-10 weeks were used. Phase of the estrus cycle was determined by assessing vaginal opening [10]. During assessing vaginal opening, female mice usually urinate, which was collected in tubes and then used immediately. Three different exposure sessions were performed. In the first session (female experience), the test mice were exposed for 5 min to a female mouse with unknown estrus cycle phase. The purpose of this session was to give the male mice social experience with females which is shown to increase female urine-induced ultrasonic vocalizations [11]. The second session (exposure to estrus female urine) was performed three days later. The male WT and $Bsn^{Dlx5/6}$cKO mice were exposed for 5 min to 20 µl of estrus female urine dropped into the corner of the open field. Since we realized that there were very low amounts of ultrasonic vocalization in response to estrus female urine but that there was much more ultrasonic vocalization in the first session, we added 1-2 days later a third session (exposure to estrus female mouse was in estrus. Between all experimental sessions, the open field was cleaned with soapy water.

#### Immunoblot analysis

For immunoblot analysis, the hippocampal tissue was homogenized in a buffer containing 0.32 M sucrose, 5 mM HEPES (pH 7.4) with protease inhibitors (cOmplete, EDTA-free protease inhibitor cocktail, Roche). The homogenate was centrifuged at 1000x g for 10 min, resulting in a nuclear fraction and other debris (pellet). Supernatant is further centrifuged at 12000x g at 4$^0$C for 20 min to obtain pellet P2, which is then resuspended in 0.32M sucrose/ 5mM Tris (pH 8.1) [12]. This fraction was used for quantitative immunoblot and proteomics analyses. Quantitative immunoblotting was performed as described previously [3] using Tris-Acetate polyacrylamide gradient gels (4-8%) for separating larger molecular weight proteins, like Bassoon, and Tris-Glycine polyacrylamide gels (5-20%) for proteins <250 kDa. Tris-Glycine gels contained 1.2% (v/v) 2,2,2-trichlorethanol (TCE), which was later used for normalization to total protein contents [13]. For Tris-acetate gels, the signal of the test protein was normalized to β-Tubulin levels. Loading was 10-20 µg protein per lane. The immunoreactive fluorescent signal (integrated density-ID) was measured using an Odyssey Infrared Scanner (LI-COR Biosciences), and the HRP/POD signal was quantified with an ECL Chemo-scanner (INTAS Science imaging instruments GmbH). Quantification of the signals was performed using Image Studio software. ID values of test proteins were normalized to the mean value of the respective WT group.

#### Proteome analysis

Proteomic experimentation and preliminary data analysis was obtained through commercial service by EMBL Proteomics Core Facility (Heidelberg) (https://www.embl.org/groups/proteomics/), employing the Liquid chromatography (LS)-MS/MS method.

*Enrichment analysis*

The false discovery rate on peptide and protein level was set to 0.01 and the fold change was set to 1.3 in order to consider a candidate to be regulated. MouseMine (www.mousemine.org) data bioinformatic tool was used to perform gene ontology (GO) enrichment analysis [14]. GraphPad Prism and Program Complex Heatmap were used to generate heatmap and UpSet plots, respectively [15].

#### Mitochondrial function assay

Primary hippocampal neurons from both the genotypes were seeded at densities of 30,000 cells per well (96-well XF cell culture microplate, Seahorse Agilent Technologies) in 200 µl of Neurobasal A medium supplemented with B27, 11 mM sodium pyruvate, 4 mM Glutamax and antibiotics (100 U/ml penicillin, 100 µg/ml streptomycin) and grown at 37°C, 5% $CO_2$. At 1 days and 3 *in vitro,* Arac (cytosine arabinoside) was added to the cells (0.6 µM each time) to reach a final concentration of 1.2 µM to suppress glial overgrowth growth.

XF sensor cartridges were hydrated with sterile water and kept overnight at 37°C in a $CO_2$-free incubator. Before experiments, they were hydrated with 200 µL of XF calibrator and kept at 37°C in a $CO_2$-free incubator for 60 minutes. Injection ports were loaded with drugs dissolved in XF assay medium. During the mitostress assay, once cells reached the desired age (14 DIV), neuronal medium was removed, neurons were washed twice with XF assay medium (Agilent XF-RPMI supplemented with 25 mM glucose, 2 mM L-glutamine, and 1 mM sodium pyruvate; pH 7.4), then replaced with 180 µL of assay medium and incubated in a

$CO_2$-free incubator at 37°C for 1 h. Oxygen consumption rate (OCR) was measured during sequential injection of 1.5 µM oligomycin, 2 µM FCCP, and 0.5 µM rotenone/antimycin A. For basal measurement, control ports were cell-free but with drug added. The cell plate and cartridge were placed on the XFe96 analyzer, and results were analyzed.

#### Statistical analysis

GraphPad Prism (GraphPad Software Inc., version 9), SigmaPlot for Windows Version 11.0 (Systat software) and JASP software (https://github.com/jasp-stats/jasp-desktop) were employed for statistical analysis in this study with respect to different experimental procedures including *in vitro,* immunohistochemical and behavioral analyses and electrophysiological recordings. In all the cases, normality test (Shapiro-Wilk test and/or Levene´s test) and equal variance test (in case of slice electrophysiology) were performed before commencing other statistical tests. One-way or two-way ANOVA (repeated measures) and paired or unpaired Students *t* tests were used. In case the data did not pass the normality test, Mann-Whitney *U* test was used. Details are indicated for each experiment in the corresponding figure captions. For the statistical comparison of gamma power, log transformed data was used to reduce variability. To determine whether probability to induce recurrent epileptiform discharges was altered, Fisher´s exact test was used. All the data are represented as means ± standard errors of the mean (SEM). Probability values of $p<0.05$ were considered as statistically significant. Sample sizes are provided in figure captions for each experiment (N: Number of mice; n: Number of cells or slices).

**Supplementary table 1.** *Detailed information on statistical analysis of input-output curves for fEPSP slopes*

| Region | Stimulation Intensity (μA) | Statistical Test | P Value |
|---|---|---|---|
| **CA1** | 5 | Mann-Whitney U Statistic=441.0 | P=0.188 |
| | 10 | Mann-Whitney U Statistic=290.0 | P=0.001 |
| | 15 | Mann-Whitney U Statistic=264.0 | P<0.001 |
| | 20 | Mann-Whitney U Statistic=272.0 | P<0.001 |
| | 30 | Mann-Whitney U Statistic=277.0 | P<0.001 |
| | 40 | Mann-Whitney U Statistic=299.0 | P=0.002 |
| | 50 | Mann-Whitney U Statistic=311.0 | P=0.003 |
| | 75 | Mann-Whitney U Statistic=344.0 | P=0.01 |
| | 100 | Mann-Whitney U Statistic=316.0 | P=0.004 |

**Supplementary table 2.** *Detailed information on statistical analysis of input-output curves for FV amplitudes*

| Region | Stimulation Intensity (μA) | Statistical Test | P Value |
|---|---|---|---|
| **CA1** | 5 | Mann-Whitney U Statistic=500.0 | P=0.718 |
| | 10 | Mann-Whitney U Statistic=387.0 | P=0.065 |
| | 15 | Mann-Whitney U Statistic=359.0 | P=0.027 |
| | 20 | Mann-Whitney U Statistic=344.0 | P=0.016 |
| | 30 | Mann-Whitney U Statistic=332.0 | P=0.01 |
| | 40 | Mann-Whitney U Statistic=320.0 | P=0.006 |
| | 50 | Mann-Whitney U Statistic=322.0 | P=0.007 |
| | 75 | Mann-Whitney U Statistic=319.0 | P=0.006 |
| | 100 | Mann-Whitney U Statistic=317.0 | P=0.006 |

**Supplementary table 3.** *Detailed information on statistical analysis of paired-pulse ratios*

| Region | Stimulus Interval (μA) | Statistical Test | P Value |
|---|---|---|---|
| **CA1** | 10 | Mann-Whitney U Statistic=383.0 | P=0.026 |
| | 25 | Mann-Whitney U Statistic=472.0 | P=0.267 |
| | 50 | Student´s two-tailed Statistic T(65)=1.408 | P=0.164 |
| | 100 | Student´s two-tailed Statistic T(65)=-1.127 | P=0.264 |
| | 250 | Mann-Whitney U Statistic=371.0 | P=0.017 |
| | 500 | Mann-Whitney U Statistic=408.0 | P=0.056 |

**Supplementary Table 4.** *Detailed information on statistical analysis of synaptic fatigue during HFS*

| Region | Stimulus # | Statistical Test | P Value |
|---|---|---|---|
| **CA1** | 2-10 | Student´s two-tailed Statistic T(18)=-2.647 | P=0.016 |
| | 11-20 | Student´s two-tailed Statistic T(18)=-1.904 | P=0.073 |
| | 21-30 | Student´s two-tailed Statistic T(18)=-1.721 | P=0.102 |
| | 31-40 | Student´s two-tailed Statistic T(18)=-1.431 | P=0.170 |
| | 41-50 | Student´s two-tailed Statistic T(18)=-1.124 | P=0.276 |
| | 51-60 | Student´s two-tailed Statistic T(18)=-1.061 | P=0.303 |
| | 61-70 | Student´s two-tailed Statistic T(18)=-0.817 | P=0.424 |
| | 71-80 | Student´s two-tailed Statistic T(18)=-0.438 | P=0.667 |
| | 81-90 | Student´s two-tailed Statistic T(18)=-0.537 | P=0.598 |
| | 91-100 | Student´s two-tailed Statistic T(18)=-0.410 | P=0.686 |

**Supplementary Table 5.** *List of candidates regulated in Bsn$^{Dlx5/6}$cKO mice in proteomics screen*

| Gene_name | UniProt Entry ID | logFC |
|---|---|---|
| S100A6 | P14069\|Q545I9 | 1.63433348 |
| TGM1 | A0A0R4J293 | 1.49225612 |
| CD44 | A2APM1\|A2APM2\|A2APM3\|A2APM4\|A2APM5\|E9QKM8\|P15379\|P15379-10\|P15379-11\|P15379-2\|P15379-3\|P15379-4\|P15379-5\|P15379-6\|P15379-7\|P15379-8\|P15379-9\|Q3U8S1\|Q80X37 | 1.32203864 |
| HSPB1 | P14602\|P14602-2\|P14602-3 | 1.1198778 |
| BDNF | A2AII2\|H9H9S8\|P21237\|Q541P3 | 1.01831625 |
| ALDH1L2 | D3Z6B9\|Q8K009 | 0.98704531 |
| APOA1 | Q00623 | 0.78364774 |
| NPTX2 | O70340 | 0.76283203 |
| TMSB4X | P20065-2 | 0.75111424 |
| UQCRH | P99028 | 0.71809876 |
| ATP5J | P97450 | 0.71701328 |
| SLC4A2 | A0A0R4J101\|A0A0R4J1K4\|A0A0R4J1K9\|P13808\|P13808-2\|P13808-3 | 0.71115658 |
| CYB5R1 | Q9DB73 | 0.69963553 |
| CAPG | Q99LB4 | 0.6906815 |
| THBS4 | B2RTL6\|Q9Z1T2 | 0.67184269 |
| INMT | P40936 | 0.65735285 |
| MSN | P26041 | 0.65426738 |
| GLIPR2 | Q9CYL5 | 0.64531203 |
| SERPINA3N | G3X8T9\|Q91WP6 | 0.63964401 |
| OCIAD2 | A0A0J9YU93\|Q9D8W7 | 0.63824716 |
| COX7A1 | A0A140LIU4\|P56392\|Q792A4 | 0.62955033 |
| TMLHE | Q91ZE0 | 0.6215568 |
| VIM | P20152\|Q5FWJ3 | 0.62132299 |
| VGF | Q0VGU4 | 0.61826805 |
| PLGRKT | Q9D3P8 | 0.61544587 |
| SLC25A42 | Q8R0Y8 | 0.61307503 |
| ANXA11 | P97384 | 0.60321026 |
| ANXA2 | P07356\|Q542G9 | 0.58979566 |
| SCG2 | Q03517\|Q4W8U9 | 0.57762739 |
| COQ3 | Q8BMS4 | 0.56896424 |
| CCDC109B | Q810S1 | 0.55908962 |
| NPTXR | E9PZM8\|F7BX42 | 0.54689719 |
| SHB | Q6PD21\|SHB_MOUSE | 0.54549563 |
| DHRS1 | Q99L04 | 0.53533524 |
| FAM210B | Q9D8B6 | 0.53213555 |
| MIF | P34884\|Q545F0 | 0.53135003 |
| SCCPDH | F6SP57\|Q8R127 | 0.53024487 |
| ATP5E | P56382\|Q545F5 | 0.52275228 |
| AHNAK2 | E9PYB0\|F7CVJ5\|F7DBB3 | 0.52116375 |

| | | |
|---|---|---|
| IMMP2L | Q8BPT6 | 0.51661349 |
| Q3UNZ8 | Q3UNZ8 | 0.50030535 |
| TAGLN | P37804 | 0.49713588 |
| COQ9 | F6SFF5\|Q8K1Z0 | 0.4914021 |
| ALDH1B1 | Q9CZS1 | 0.49130133 |
| TIMM10B | Q9WV96 | 0.49108867 |
| VWA5A | Q99KC8 | 0.47898229 |
| CFH | A0A0A6YWP4\|D6RGQ0\|E9Q8H9\|E9Q8I0\|P06909 | 0.47838237 |
| SV2C | A2RSH4\|Q69ZS6 | 0.47817032 |
| GM9774\|ADRM1 | A0A0A6YVU8\|Q9JKV1 | 0.47761667 |
| ACAN | Q61282 | 0.47172917 |
| ME3 | Q8BMF3 | 0.47022388 |
| PGAM2 | O70250\|Q5NCI4 | 0.46913053 |
| CAV1 | D3Z0J2\|D3Z148\|H3BKG0\|P49817\|P49817-2 | 0.46840253 |
| PDK1 | Q8BFP9 | 0.46750156 |
| CD9 | P40240 | 0.46456862 |
| SLC14A1 | Q8VHL0\|Q8VHL0-2 | 0.4636915 |
| ADCK3 | Q60936 | 0.45883948 |
| MTCH2 | Q791V5 | 0.45737868 |
| SLC25A32 | Q8BMG8 | 0.45494947 |
| CYB5R3 | Q9DCN2\|Q9DCN2-2 | 0.45414362 |
| TPP1 | O89023 | 0.45066441 |
| DYNLL2 | Q9D0M5 | 0.44469292 |
| GFAP | P03995\|P03995-2 | 0.44345431 |
| PDLIM4 | P70271 | 0.44124583 |
| COA3 | Q9D2R6 | 0.44078765 |
| GPT2 | Q8BGT5 | 0.44058785 |
| SLC25A20 | Q9Z2Z6 | 0.43933555 |
| MRPS16 | Q9CPX7 | 0.43833938 |
| CPQ | Q9WVJ3\|Q9WVJ3-2 | 0.43795227 |
| TNC | Q80YX1\|Q80YX1-2 | 0.43667853 |
| NDUFA1 | O35683\|Q545K0 | 0.43392256 |
| ACAT1 | Q8QZT1 | 0.43182797 |
| SORCS1 | E9PYT6\|E9QQ02\|E9QQ63\|Q9JLC4\|Q9JLC4-2\|Q9JLC4-3\|Q9JLC4-4 | 0.43182627 |
| MIC13 | Q8R404 | 0.43093435 |
| SLC25A51 | A2AKW0\|Q5HZI9 | 0.42882028 |
| FAM129B | Q8R1F1\|NIBA2_MOUSE | 0.42627315 |
| CD109 | A6MDD3\|Q8R422 | 0.42590738 |
| SLC16A1 | P53986\|Q544N9 | 0.42499394 |
| RIN1 | Q921Q7 | 0.42137203 |
| MLC1 | E9QP87\|Q8VHK5 | 0.41958998 |
| RGS14 | P97492 | 0.41893737 |
| ECE1 | Q4PZA2\|Q4PZA2-2\|Q4PZA2-3\|Q4PZA2-4 | 0.41723957 |
| USP30 | A0A0G2JDF7\|Q3UN04 | 0.41383957 |

| | | |
|---|---|---|
| UQCRB | Q9CQB4\|Q9D855 | 0.41369848 |
| Q8R1Y2 | Q8R1Y2 | 0.41309443 |
| 2310061I04RIK | B8JJ66\|B8JJ69 | 0.41095714 |
| AGPAT5 | F8WGD9\|Q9D1E8 | 0.41000176 |
| NDUFC2 | Q9CQ54 | 0.40657253 |
| SCRG1 | O88745\|Q543T5 | 0.40646696 |
| COX6A1 | P43024\|Q9DCW5 | 0.40411363 |
| APOE | P08226\|Q3TXU4 | 0.40266056 |
| SYN3 | Q8JZP2 | 0.40019368 |
| PITRM1 | Q8K411\|Q8K411-2 | 0.39822848 |
| PLPP4 | Q0VBU9 | 0.39765726 |
| MRPS12 | O35680 | 0.39585839 |
| SDCBP | O08992\|Q3TMX0 | 0.39460147 |
| STAMBP | Q9CQ26 | 0.39421858 |
| B2M | P01887 | 0.39372285 |
| GM9755\|TUFM | D3YVN7\|Q8BFR5 | 0.39365674 |
| COQ6 | Q8R1S0 | 0.39270155 |
| TAGLN2 | Q9WVA4 | 0.39003756 |
| FLNA | B7FAU9\|Q8BTM8 | 0.38814368 |
| ADCK5 | E0CXW2\|E9PUK2\|Q80V03\|Q80V03-2 | 0.38755185 |
| PLAT | P11214 | 0.38659763 |
| GM20498 | E9PVN6 | 0.38533803 |
| MARCH5 | A2RTC8\|Q3KNM2 | 0.38289749 |
| FECH | P22315\|Q544X6 | 0.38153326 |
| SRSF1 | H7BX95\|Q6PDM2 | 0.38137062 |
| NPTX1 | A2ACL9\|Q62443 | 0.38044815 |
| ABCB10 | Q9JI39 | 0.37997837 |
| NAP1L1 | E9PW66\|P28656 | -0.3796496 |
| CORO2A | B1AVH4\|B1AVH5\|Q8C0P5 | -0.3817203 |
| CALB1 | P12658 | -0.382793 |
| PPP1R9A | H3BJD6\|H3BL28\|Q7TN74 | -0.3834233 |
| GRM3 | Q9QYS2 | -0.3836343 |
| MAGI1 | A0A0N4SUZ0\|Q6RHR9\|Q6RHR9-3 | -0.3880147 |
| HAPLN2 | B2RRU7\|Q9ESM3 | -0.3892685 |
| CRHR1 | P35347\|Q3ZAT0 | -0.3910045 |
| DHX9 | E9QNN1 | -0.3926445 |
| GRB14 | A2ASX2\|Q9JLM9 | -0.4007612 |
| DCXR | Q91X52 | -0.4009065 |
| LEMD3 | D3YU56\|E9QP59\|Q9WU40\|Q9WU40-2 | -0.4082945 |
| MLF1 | Q3V042\|Q8JZS2\|Q9QWV4 | -0.4090285 |
| NGEF | E9QK62\|Q8CHT1\|Q8CHT1-2 | -0.4092437 |
| MTR | A6H5Y3 | -0.4111899 |
| PTGDS | O09114\|O09114-2 | -0.4122484 |
| SMARCAL1 | Q8BJL0\|Q8BJL0-2 | -0.4183687 |
| CLTA | B1AWD8\|B1AWD9\|B1AWE0\|O08585\|Q6PFA2 | -0.4184321 |

| | | |
|---|---|---|
| ADCY1 | O88444 | -0.4189449 |
| VAC14 | Q80WQ2 | -0.4202929 |
| OCLN | B2RS24\|E0CZ73\|Q61146 | -0.425553 |
| TMEM163 | Q8C996 | -0.4259515 |
| RASAL1 | Q9Z268 | -0.4269019 |
| MYO5B | G3X9Y9\|G5E8G6\|P21271\|P21271-3 | -0.4300487 |
| TBL1X | Q9QXE7 | -0.4300901 |
| CA4 | Q64444 | -0.4313168 |
| MFGE8 | P21956\|P21956-2 | -0.4315468 |
| CLTB | Q6IRU5 | -0.4334293 |
| HMGCS1 | Q8JZK9 | -0.4341628 |
| GRIK3 | B1AS29\|GRIK3_MOUSE | -0.4379778 |
| ATP2B4 | E9Q828\|F7AAP4 | -0.4385313 |
| CPSF7 | Q8BTV2\|Q8BTV2-2 | -0.4411179 |
| GYS1 | Q9Z1E4 | -0.4473139 |
| VAT1L | Q80TB8 | -0.4567356 |
| EPB42 | P49222\|Q3UV95 | -0.4578426 |
| SLC17A6 | A0A0R4J0A6\|Q8BLE7 | -0.4602493 |
| SAFB | D3YXK2\|S4R1M2 | -0.4611992 |
| ITPKA | Q8R071 | -0.4636828 |
| PYGM | Q9WUB3 | -0.4706485 |
| RGS12 | D3Z0G5\|D3Z0G6 | -0.4733285 |
| GM28062\|HNRNPU | G3XA10\|Q8VEK3 | -0.4749919 |
| ILF3 | Q9Z1X4\|Q9Z1X4-3 | -0.4790583 |
| SLC13A5 | A0A140LIC4\|A0A140LIR1\|Q5NBV0\|Q67BT3 | -0.4836439 |
| NOVA1 | Q9JKN6 | -0.4847645 |
| DSP | E9PZW0\|E9Q557 | -0.4943233 |
| C530008M17RIK\|KIAA1211 | E9Q3Y7\|E9Q5L4\|Q5PR69\|Q5PR69-2 | -0.5030863 |
| CPNE5 | Q8JZW4 | -0.5038777 |
| NPY1R | Q04573 | -0.515573 |
| HIST1H1B | P43276 | -0.5394878 |
| LBR | A0A0A6YW01\|A0A0A6YXT6\|A0A0A6YXW3\|Q3U9G9 | -0.5593285 |
| ZNFX1 | Q8R151 | -0.5690865 |
| SYNJ2 | D3YWM9\|D3YZB2\|D3YZB3\|D3YZB4\|D3Z6E7\|E9Q4P5\|F8WHD8\|Q9D2G5\|Q9D2G5-2\|Q9D2G5-3\|Q9D2G5-4\|Q9D2G5-5\|Q9D2G5-6 | -0.5780178 |
| FAM81A | Q3UXZ6 | -0.5846486 |
| CD34 | Q64314\|Q64314-2 | -0.6163898 |
| FDFT1 | P53798 | -0.6565687 |
| H2AFV\|H2AFZ | B2RVP5\|P0C0S6\|Q3THW5\|Q3UA95 | -0.6764822 |
| DYNLRB1 | P62627 | -0.702802 |
| CHN2 | Q3V2R3 | -0.7133472 |
| RAD23B | P54728 | -0.7150382 |
| SLC4A1 | P04919\|P04919-2 | -0.7225617 |
| IGKC\|P01837 | A0A075B5P2\|P01837 | -0.7344202 |
| SF3B1 | G5E866\|Q99NB9 | -0.8249095 |

| | | |
|---|---|---|
| CENPF | E9Q3P4 | -0.9321289 |
| BBS4 | A6H669\|Q8C1Z7 | -1.0478298 |
| LMNA | P48678\|P48678-2 | -1.0528096 |
| LMNB1 | P14733 | -1.1901761 |
| RANBP2 | Q9ERU9 | -1.1988292 |
| IGHV1 | A0A075B5Y4\|A0A0A6YVW3 | -1.2665313 |
| LMNB2 | P21619 | -1.3632291 |

**Supplementary Table 6.** *List of GO-cellular component enrichment terms in Bsn*$^{Dlx5/6}$*cKO*

| GO Term | GO | p-Value (Holm-Bonferroni p 0.05) |
|---|---|---|
| organelle inner membrane | GO:0019866 | 7.956007998397744e-16 |
| organelle envelope/envelope | GO:0031967/GO:0031975 | 1.4075506799775919e-15 |
| mitochondrial membrane | GO:0031966 | 1.900331980759324e-14 |
| mitochondrial inner membrane | GO:0005743 | 2.96061804401796e-14 |
| mitochondrion | GO:0005739 | 1.6597382604289894e-13 |
| mitochondrial envelope | GO:0005740 | 1.8238814526723002e-13 |
| organelle membrane | GO:0031090 | 2.0337542912181602e-12 |
| cytoplasm | GO:0005737 | 4.747526366593665e-10 |
| organelle | GO:0043226 | 1.2786774814289343e-7 |
| intracellular organelle | GO:0043229 | 1.9122783519507765e-7 |
| intracellular anatomical structure | GO:0005622 | 0.0000017573062373257909 |
| intracellular membrane-bounded organelle | GO:0043231 | 0.000005870088763886164 |
| inner mitochondrial membrane protein complex | GO:0098800 | 0.00004103304322463525 |
| membrane-bounded organelle | GO:0043227 | 0.000042293519968352415 |
| mitochondrial protein-containing complex | GO:0098798 | 0.00008485191304635309 |
| plasma membrane region | GO:0098590 | 0.00019632597434898858 |
| cell junction | GO:0030054 | 0.00037414313981514526 |
| glutamatergic synapse | GO:0098978 | 0.0008479652492709834 |
| synapse | GO:0045202 | 0.005262538795330125 |
| basal plasma membrane | GO:0009925 | 0.010954781724491401 |
| basolateral plasma membrane | GO:0016323 | 0.017728817710839763 |
| nuclear lamina | GO:0005652 | 0.01982802486540773 |
| basal part of cell | GO:0045178 | 0.027798790368714882 |
| cell projection | GO:0042995 | 0.03143404823055427 |
| lamin filament | GO:0005638 | 0.04158274717516748 |
| ubiquinone biosynthesis complex | GO:0110142 | 0.04158274717516748 |
| mitochondrial respirasome | GO:0005746 | 0.042269273422046116 |

**Supplementary Table 7.** *List of overlapped candidates among top five GO-cellular component terms (1-presence; 0-absence)*

| ID | Organelle inner membrane | Organelle envelope | Mitochondrial membrane | Mitochondrial Innermembrane | Mitochondria |
|---|---|---|---|---|---|
| 2310061I04Rik | 0 | 0 | 0 | 0 | 1 |
| Abcb10 | 1 | 1 | 1 | 1 | 1 |
| Acat1 | 1 | 1 | 1 | 1 | 1 |
| Adck5 | 0 | 0 | 0 | 0 | 1 |
| Agpat5 | 0 | 1 | 0 | 0 | 1 |
| Aldh1b1 | 0 | 0 | 0 | 0 | 1 |
| Aldh1l2 | 0 | 0 | 0 | 0 | 1 |
| Anxa11 | 0 | 1 | 0 | 0 | 0 |
| Apoe | 0 | 1 | 0 | 0 | 0 |
| Atp5e | 1 | 1 | 1 | 1 | 1 |
| Atp5j | 1 | 1 | 1 | 1 | 1 |
| Bdnf | 1 | 1 | 1 | 1 | 1 |
| Cav1 | 0 | 0 | 0 | 0 | 1 |
| Cenpf | 0 | 1 | 0 | 0 | 0 |
| Cfh | 0 | 0 | 0 | 0 | 1 |
| Coa3 | 1 | 1 | 1 | 1 | 1 |
| Coq3 | 1 | 1 | 1 | 1 | 1 |
| Coq6 | 1 | 1 | 1 | 1 | 1 |
| Coq9 | 1 | 1 | 1 | 1 | 1 |
| Cox6a1 | 1 | 1 | 1 | 1 | 1 |
| Cox7a1 | 1 | 1 | 1 | 1 | 1 |
| Cyb5r1 | 0 | 0 | 0 | 0 | 1 |
| Cyb5r3 | 1 | 1 | 1 | 1 | 1 |
| Dhrs1 | 1 | 1 | 1 | 1 | 1 |
| Fam210b | 0 | 1 | 1 | 0 | 1 |
| Fech | 1 | 1 | 1 | 1 | 1 |
| Gpt2 | 0 | 0 | 0 | 0 | 1 |
| Ilf3 | 0 | 0 | 0 | 0 | 1 |
| Immp2l | 1 | 1 | 1 | 1 | 1 |
| Lbr | 1 | 1 | 0 | 0 | 0 |
| Lemd3 | 1 | 1 | 0 | 0 | 0 |
| Lmna | 0 | 1 | 0 | 0 | 0 |
| Lmnb1 | 1 | 1 | 0 | 0 | 0 |
| Lmnb2 | 0 | 1 | 0 | 0 | 0 |
| Marchf5 | 0 | 1 | 1 | 0 | 1 |
| Me3 | 0 | 0 | 0 | 0 | 1 |
| Micos13 | 1 | 1 | 1 | 1 | 1 |
| Mrps12 | 1 | 1 | 1 | 1 | 1 |
| Mrps16 | 1 | 1 | 1 | 1 | 1 |
| Mtch2 | 1 | 1 | 1 | 1 | 1 |

| | | | | | |
|---|---|---|---|---|---|
| Ndufa1 | 1 | 1 | 1 | 1 | 1 |
| Ndufc2 | 1 | 1 | 1 | 1 | 1 |
| Nptx1 | 0 | 0 | 0 | 0 | 1 |
| Ociad2 | 1 | 1 | 1 | 1 | 1 |
| Pdk1 | 0 | 0 | 0 | 0 | 1 |
| Pitrm1 | 0 | 0 | 0 | 0 | 1 |
| Plgrkt | 0 | 0 | 0 | 0 | 1 |
| Ranbp2 | 0 | 1 | 0 | 0 | 1 |
| S100a6 | 0 | 1 | 0 | 0 | 0 |
| Sccpdh | 0 | 0 | 0 | 0 | 1 |
| Sdcbp | 0 | 1 | 0 | 0 | 0 |
| Slc16a1 | 0 | 0 | 0 | 0 | 1 |
| Slc25a20 | 1 | 1 | 1 | 1 | 1 |
| Slc25a32 | 1 | 1 | 1 | 1 | 1 |
| Slc25a42 | 1 | 1 | 1 | 1 | 1 |
| Slc25a51 | 1 | 1 | 1 | 1 | 1 |
| Synj2 | 0 | 1 | 1 | 0 | 1 |
| Timm10b | 1 | 1 | 1 | 1 | 1 |
| Tmlhe | 0 | 0 | 0 | 0 | 1 |
| Tpp1 | 0 | 0 | 0 | 0 | 1 |
| Uqcrb | 1 | 1 | 1 | 1 | 1 |
| Uqcrh | 1 | 1 | 1 | 1 | 1 |
| Usp30 | 0 | 1 | 1 | 0 | 1 |
| Znfx1 | 0 | 1 | 1 | 0 | 1 |

**Supplementary Table 8.** *List of GO-biological processes enrichment terms in Bsn*$^{Dlx5/6}$*cKO*

| GO Term | GO | p-Value (Holm-Bonferroni p 0.05) |
|---|---|---|
| oxidative phosphorylation | GO:0006119 | 0.00012126290577767516 |
| generation of precursor metabolites and energy | GO:0006091 | 0.0001815595867946688 |
| system development | GO:0048731 | 0.0014631052426385017 |
| ATP metabolic process | GO:0046034 | 0.002517673073232617 |
| sterol biosynthetic process | GO:0016126 | 0.0032400912602044865 |
| aerobic respiration | GO:0009060 | 0.004154280081338271 |
| localization | GO:0051179 | 0.008041473687234539 |
| blood coagulation | GO:0007596 | 0.00917783329711572 |
| hemostasis | GO:0007599 | 0.011058523343361153 |
| coagulation | GO:0050817 | 0.011058523343361153 |
| leukocyte aggregation | GO:0070486 | 0.011486309798819344 |
| electron transport chain | GO:0022900 | 0.01230371336880541 |
| cellular respiration | GO:0045333 | 0.014142090138730983 |
| energy derivation by oxidation of organic compounds | GO:0015980 | 0.01786375162498987 |
| cholesterol biosynthetic process | GO:0006695 | 0.024163883691910117 |
| secondary alcohol biosynthetic process | GO:1902653 | 0.024163883691910117 |
| transport | GO:0006810 | 0.03429383859722221 |
| small molecule metabolic process | GO:0044281 | 0.04240976038977984 |

**Supplementary Table 9.** *List of overlapped candidates among top five GO-biological processes terms (1-presence; 0-absence)*

| ID | Oxidative phosphorylation | Precursor metabolites | System development | ATP metabolic process | Sterol biosynthesis |
|---|---|---|---|---|---|
| Abcb10 | 0 | 0 | 1 | 0 | 0 |
| Acan | 0 | 0 | 1 | 0 | 0 |
| Acat1 | 0 | 1 | 0 | 0 | 0 |
| Adcy1 | 0 | 0 | 1 | 0 | 0 |
| Agpat5 | 0 | 0 | 1 | 0 | 0 |
| Aldh1l2 | 0 | 1 | 0 | 0 | 0 |
| Anxa2 | 0 | 0 | 1 | 0 | 0 |
| Apoa1 | 0 | 0 | 1 | 0 | 1 |
| Apoe | 0 | 0 | 1 | 0 | 1 |
| Atp2b4 | 0 | 0 | 1 | 0 | 0 |
| Atp5e | 1 | 1 | 0 | 1 | 0 |
| Atp5j | 1 | 1 | 0 | 1 | 0 |
| B2m | 0 | 0 | 1 | 0 | 0 |
| Bbs4 | 0 | 0 | 1 | 0 | 0 |
| Bdnf | 1 | 1 | 1 | 1 | 0 |
| Capg | 0 | 0 | 1 | 0 | 0 |
| Cav1 | 0 | 0 | 1 | 0 | 0 |
| Cd109 | 0 | 0 | 1 | 0 | 0 |
| Cd34 | 0 | 0 | 1 | 0 | 0 |
| Cd44 | 0 | 0 | 1 | 0 | 0 |
| Cd9 | 0 | 0 | 1 | 0 | 0 |
| Cenpf | 0 | 0 | 1 | 0 | 0 |
| Cfh | 0 | 0 | 1 | 1 | 0 |
| Coq9 | 1 | 1 | 0 | 1 | 0 |
| Cox6a1 | 1 | 1 | 0 | 1 | 0 |
| Cox7a1 | 1 | 1 | 0 | 1 | 0 |
| Crhr1 | 0 | 0 | 1 | 0 | 0 |
| Cyb5r1 | 0 | 0 | 0 | 0 | 1 |
| Cyb5r3 | 0 | 0 | 0 | 0 | 1 |
| Dsp | 0 | 0 | 1 | 0 | 0 |
| Ece1 | 0 | 0 | 1 | 0 | 0 |
| Epb42 | 0 | 0 | 1 | 0 | 0 |
| Fam210b | 0 | 0 | 1 | 0 | 0 |
| Fdft1 | 0 | 0 | 0 | 0 | 1 |
| Fech | 0 | 0 | 1 | 0 | 0 |
| Flna | 0 | 0 | 1 | 0 | 0 |
| Gfap | 0 | 0 | 1 | 0 | 0 |
| Gys1 | 0 | 1 | 1 | 0 | 0 |
| Hapln2 | 0 | 0 | 1 | 0 | 0 |
| Hmgcs1 | 0 | 0 | 0 | 0 | 1 |
| Hspb1 | 0 | 0 | 1 | 0 | 0 |

| | | | | | |
|---|---|---|---|---|---|
| Immp2l | 0 | 1 | 1 | 0 | 0 |
| Itpka | 0 | 0 | 1 | 0 | 0 |
| Lbr | 0 | 0 | 1 | 0 | 1 |
| Lemd3 | 0 | 0 | 1 | 0 | 0 |
| Lmna | 0 | 0 | 1 | 0 | 0 |
| Mfge8 | 0 | 0 | 1 | 0 | 0 |
| Mlf1 | 0 | 0 | 1 | 0 | 0 |
| Msn | 0 | 0 | 1 | 0 | 0 |
| Mtch2 | 1 | 1 | 0 | 1 | 0 |
| Mtr | 0 | 0 | 1 | 0 | 0 |
| Myo5b | 0 | 0 | 1 | 0 | 0 |
| Nap1l1 | 0 | 0 | 1 | 0 | 0 |
| Ndufa1 | 1 | 1 | 0 | 1 | 0 |
| Ndufc2 | 1 | 1 | 0 | 1 | 0 |
| Ngef | 0 | 0 | 1 | 0 | 0 |
| Nova1 | 0 | 0 | 1 | 0 | 0 |
| Nptx1 | 0 | 0 | 1 | 0 | 0 |
| Nptxr | 0 | 0 | 1 | 0 | 0 |
| Npy1r | 0 | 0 | 1 | 0 | 0 |
| Pdlim4 | 0 | 0 | 1 | 0 | 0 |
| Pgam2 | 0 | 1 | 0 | 1 | 0 |
| Ppp1r9a | 0 | 0 | 1 | 0 | 0 |
| Pygm | 0 | 1 | 0 | 0 | 0 |
| Rgs14 | 0 | 0 | 1 | 0 | 0 |
| Scg2 | 0 | 0 | 1 | 0 | 0 |
| Sdcbp | 0 | 0 | 1 | 0 | 0 |
| Shb | 0 | 0 | 1 | 0 | 0 |
| Slc17a6 | 0 | 0 | 1 | 0 | 0 |
| Slc4a1 | 0 | 1 | 1 | 1 | 0 |
| Slc4a2 | 0 | 0 | 1 | 0 | 0 |
| Synj2 | 0 | 0 | 1 | 0 | 0 |
| Tgm1 | 0 | 0 | 1 | 0 | 0 |
| Thbs4 | 0 | 0 | 1 | 0 | 0 |
| Tnc | 0 | 0 | 1 | 0 | 0 |
| Tpp1 | 0 | 0 | 1 | 0 | 0 |
| Uqcrb | 1 | 1 | 0 | 1 | 0 |
| Uqcrh | 1 | 1 | 0 | 1 | 0 |
| Vgf | 0 | 1 | 1 | 0 | 0 |
| Vim | 0 | 0 | 1 | 0 | 0 |
